# Object manipulation and affordance learning in *Drosophila*

**DOI:** 10.64898/2026.04.28.721021

**Authors:** Matthias Durrieu, Dominic Dall’Osto, Thomas Ka Chung Lam, Victor Lobato-Rios, Pavan Ramdya

## Abstract

Animals must manipulate objects to perform tasks like pushing away debris while navigating over complex, natural terrain. For previously unseen objects, efficient manipulation requires that their affordances—the possible actions one can perform upon them—first be learned through experience. However, the behavioral and neural mechanisms underlying the learning of object affordances remain largely unknown. To address this gap, we show that adult flies can learn to push small spherical objects without being given any explicit reward. To do this, flies appear to learn the ball’s pushability affordance: pushing is delayed when animals are first exposed to an immobile ball, and manipulating one ball accelerates pushing of a second one in a new context. Behavioral quantification of a largescale neural silencing screen reveals that specific visual projection neurons and olfactory sensory neurons regulate initial reactions to the object while dopaminergic neurons and circuits in the mushroom bodies, a center for associative learning and memory in insects, are critical for learning object affordances. These findings open the door to a mechanistic understanding of object manipulation and affordance learning.

## Introduction

Object manipulation rests at the core of exploring and exploiting the natural world. Many species manipulate objects ^1^ including humans ^2^, primates ^3^, rodents ^4,5^, and crows ^6^ as well as insects (e.g., ants carry food in their mandibles ^7^, and both bees ^8^ and dung beetles ^9^ roll balls). To accomplish this task, animals must continuously identify which objects, novel or familiar, can be manipulated and how. For example, a morsel of food might be stuck to the ground, making it futile for a foraging ant to attempt to bring it to the nest. The possible actions one can perform upon an object are known as its ‘affordances’ ^10^. These can be learned and retained, facilitating engagements with similar objects in the future. Because affordances serve as efficient action-centered models of the world, it has made them a useful framework for manipulation learning in robotics ^11,12^.

In insects, object affordances have been studied extensively at the behavioral level, including in bumble bees ^13^ and ants ^14^. However, the underlying neural mechanisms remain unknown. This gap is largely due to the fact that animal models typically used to study object affordances are not amenable to precise neural circuit-level investigation. By contrast, the fly, *Drosophila melanogaster*, gives experimenters access to powerful genetic tools ^15^ for silencing or activating neurons, the ability to explore connectivity graphs of the entire nervous system ^16,17^, and techniques for recording the activity of neurons in the brain ^18,19^ and motor system ^20^ during tethered behavior.

Until now it has not been clear to what extent flies can manipulate objects and learn their affordances. Prior studies have shown that flies tend to approach and mount objects that are immobile ^21^ and of specific shapes ^22^, but these works focused on object preferences and spatial localization rather than object manipulation. Therefore, it remains unknown: (i) whether flies manipulate small objects with their limbs, (ii) whether they can learn to manipulate objects more effectively with experience, (iii) whether manipulating an object involves learning its affordances, and (iv) which brain regions and neural circuits are involved **(Fig. 1a)**.

**Fig. 1:**
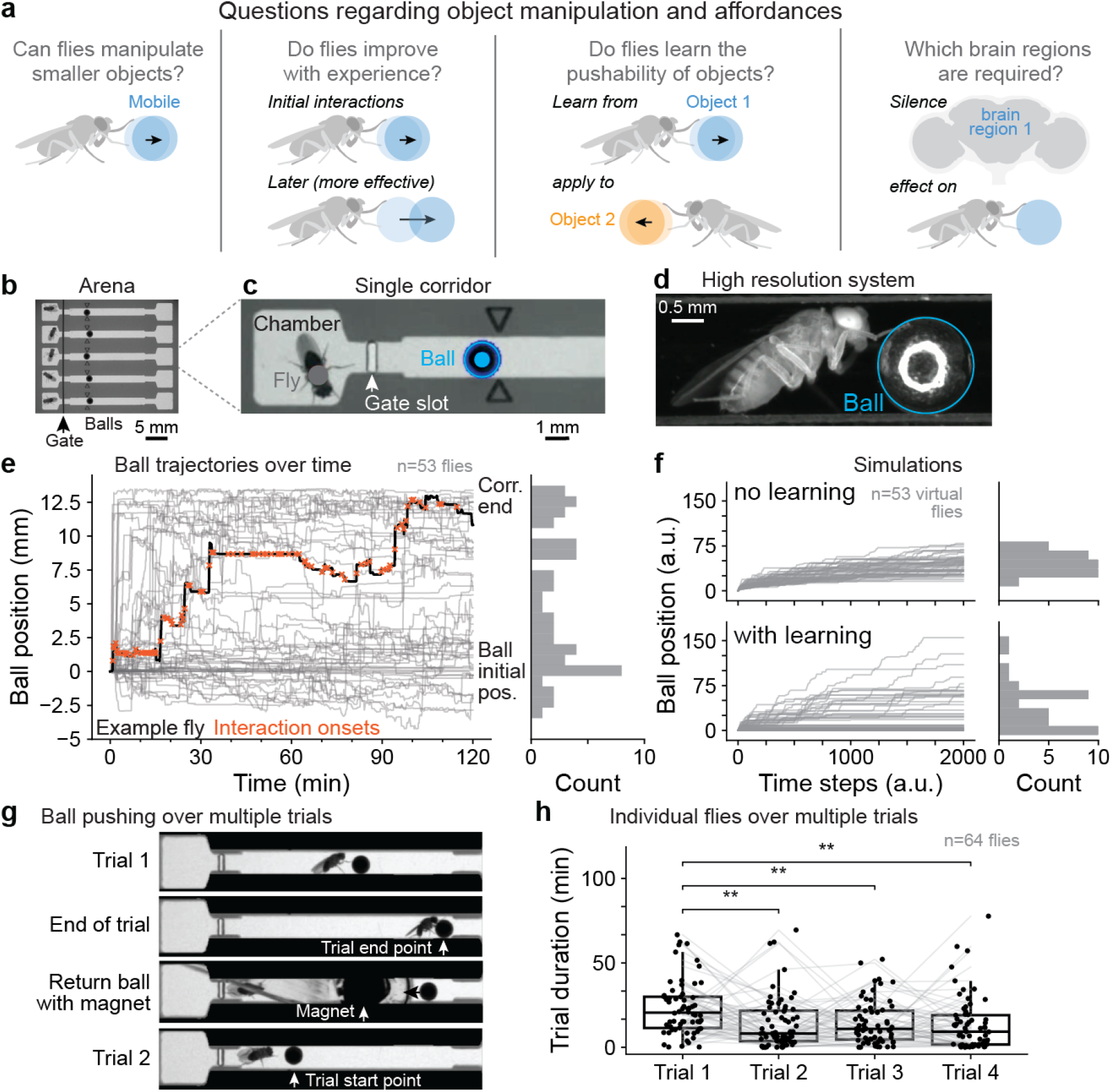
Flies manipulate objects in the absence of an extrinsic reward. **(a)** Our study addresses four questions: whether flies manipulate smaller objects with their legs, whether this manipulation improves with experience, if they learn object-specific affordances, and which neural circuits are required for this learning. **(b)** An experimental arena consisting of six narrow corridors. Each corridor is book-ended by larger starting chambers. The fly begins in the left-most chamber and is initially restricted from entering the corridor by a gate. **(c)** Zoomed-in view of one chamber and corridor. The positions of the fly’s thorax (gray dot) and the ball (blue circle and dot) are tracked using 2D pose estimation software. Two engraved black arrowheads facilitate consistent initial positioning of the ball. **(d)** High-resolution image of a fly contacting the ball (blue circle). **(e)** Ball trajectories relative to the initial starting position over two hours (n = 53 flies). Each animal’s trajectory is shown in gray. One sample trajectory (black line) overlaid with the onsets of ball contact events (orange crosses). A histogram (right) shows the distribution of final ball positions for all flies. **(f)** Computational simulations of ball manipulation behaviors (n = 53 virtual flies) either (top) without learning, using random forward and backward walking, or (bottom) adding a learning rule in which successful ball pushes increase the probability of subsequent interactions while unsuccessful pushes decrease the probability. Ball positions over time are plotted in arbitrary units over arbitrary time steps. Histograms (right) show the distribution of final ball positions for each condition. **(g)** Multitrial ball pushing experiment. Each time a fly would bring the ball to the end of the corridor, the ball would be displaced back to the initial position, marking the start of a new trial. This continued for 2 h. **(h)** Completion time (in minutes) of successive trials for individual flies that completed ≥ 4 trials. (n = 64 flies). Boxplots show the median (center line), interquartile range (IQR; box edges), 1.5 × IQR (whiskers), and individual data points (dots). Gray lines connect measurements from the same individuals across trials. Trial durations were compared using a Friedman test (*χ*^2^ = 23.0, *p* = 4.1 × 10^*−*5^) followed by pairwise post-hoc Wilcoxon signed-rank tests (FDR-corrected, *α* = 0.05). Significance levels are indicated as: ns, *p* ≥ 0.05; *, *p <* 0.05; **, *p <* 0.01; ***, *p <* 0.001.

To address these questions, we developed a behavioral assay in which flies are placed in a corridor with a smaller ball. We find that flies initially probe the ball with their forelegs and retreat but, in time, begin to push the ball down the corridor. They will perform ball pushing over multiple trials, even in the absence of an extrinsic reward like food. We could envision three potential learning mechanisms underlying these observations: (i) sensory habituation or increased motivation may modify how frequently animals interact with the ball, (ii) motor learning may increase the efficiency of fly-ball manipulations, and/or (iii) affordance learning of ball pushability may unlock pushing behaviors. Our results suggest that initial retreat reactions to the ball are mediated by vision. Furthermore, motor programs to push the ball can be performed almost immediately, implying that substantial motor learning is not required. However, flies do appear to learn the pushability affordance of the ball: if flies were previously exposed to an immobile ball, they are delayed in pushing it when it becomes mobile. Additionally, pushing one ball accelerates pushing a second ball in a different context. To investigate the brain regions involved in this behavior, we performed a high-throughput neural silencing screen of 212 driver lines targeting sparse sets of neurons across brain regions. Our neural silencing results support a framework in which two mechanistically distinct processes co-regulate ball manipulation: (i) vision- and olfaction-mediated initial reactions to these novel objects, and (ii) dopamine- and mushroom body-dependent learning of the ball’s pushability affordance.

## Results

### Flies manipulate objects in the absence of an extrinsic reward

To investigate affordance learning in *Drosophila*, we developed an experimental paradigm in which female flies manipulate a previously unseen spherical object. Specifically, we designed an arena consisting of narrow linear corridors book-ended by larger chambers in which flies are initially placed **(Fig. 1b-c)**. When a gate is opened, flies can enter the corridor and encounter a 1.5 mm diameter stainless steel ball with which they can subsequently interact **(Fig. 1d)**. For each experiment we tracked ^23^ movements of flies and balls over two hours **(Video 1)**. By pooling animals from nine such experiments, we can observe that flies push and pull the ball in discrete contact events **(Fig. 1e)** and that across animals there is considerable heterogeneity in ball pushing efficacy (the distance the ball is pushed over two hours); some push the ball to nearly the end of the corridor, whereas others fail to do so or even pull the ball back upon themselves. This results in a polarized distribution of final ball positions.

This distribution suggests that a non-random process, possibly learning, underlies the dynamics of ball manipulation. To more quantitatively explore this possibility, we simulated ball manipulation using a random walk model in which the probability of successfully pushing the ball either remains static, or is reinforced following successful pushes and suppressed following unsuccessful ones. Using a static model without learning, simulations resulted in a relatively normal spread of final ball positions **(Fig. 1f, top)**. By contrast, introducing a learning rule produced a more polarized distribution in which one group of simulated agents pushed effectively and another stalled near the start **(Fig. 1f, bottom)**. These simulation results predict that real fly behaviors do not result from a static probability policy, but that animals may learn to push more or less persistently depending on the outcome of interactions with the ball.

To estimate the time course of this potential learning process, we measured the time taken for real individual animals to push the ball to the end of the corridor across multiple trials. To do this, when a fly brought the ball near the corridor end we used a magnet to return the ball to the start position with the fly retreating as well **(Fig. 1g)**. We analyzed data from 64 (out of 161) wild-type animals that had completed at least four trials and observed a significant reduction in the time it took to bring the ball to the end of the corridor between trials 1 and 2, but no further improvement thereafter **(Fig. 1h)**. This suggests that during the first trial flies may undergo a relatively discrete transition from inefficient to efficacious ball manipulation.

### Flies learn the affordance of object pushability

Improvements in ball-pushing proficiency could result from: (i) motor learning of new object manipulation movements or fine-tuning of existing movements; (ii) affordance learning of which pre-existing movements to apply to the object; and/or (iii) general increases in the fly’s motivation to engage with these novel objects. To distinguish between these possibilities we began by visualizing fly-ball interactions at higher spatial resolution **(Fig. 1d)**. We found that flies tend to initially make brief probing contacts with the ball and often retreat afterwards **(Video 2)**. Over time flies become less likely to retreat and, ultimately, start to push the ball using strategies like rearing up and pushing with their forelegs while bracing their rear and middle legs against the corridor walls **(Video 3)**.

To quantify these observations, we manually classified ball contact events until a first ‘major’ contact event (a push of 1.2 mm or more). We classified events as either involving contact with the head, a single foreleg, both forelegs, or rearing up with both forelegs on the ball simultaneously **(Video 4)(Extended Data Fig. 1a)**. One relatively trivial explanation for ball pushing might be that flies simply walk forwards, ignore the ball and hit it with their heads. However, we found that head pushing was exceedingly rare (n = 6/692 minor and 0/38 major events). By contrast, pushing with the legs was very common. Single-leg contacts were most frequent (n = 475/692 minor and 10/38 major events), followed by contact with both forelegs (n = 159/692 minor and 5/38 major events). Finally, rearing up on the ball and pushing it with both forelegs was less common (n = 52/692 minor and 23/38 major events), but the most effective movement for pushing the ball the farthest **(Extended Data Fig. 1b)**.

Notably, we found that for some animals (n = 13/42) the first major push was achieved using a new behavioral strategy, rather than an improved version of a previously attempted one. In most cases this behavior was rearing up with their front legs on the ball while bracing with their hind legs (12/13 major pushes) **(Extended Data Fig. 1a, blue stars)**. That flies can use this movement for effective pushing on their first attempt suggests that it may be an innate motor program. Because we do not find evidence that flies need to progressively fine-tune their motor program for successful ball pushing, we conclude that improvements in pushing over time are likely due to (i) increased overall engagement with the ball and/or (ii) learning when and how to apply pre-existing motor programs to this novel object (i.e., learning the ball’s action possibilities or affordances).

To distinguish between increased motivation to engage with the ball versus affordance learning, we performed a new experiment in which we asked whether direct experience with an immobile object reduces interactions with it **(Fig. 2a, left)**. This would be inconsistent with increased engagement explaining improvements in ball pushing over time. To test this possibility, for the first hour of the experiment we fixed the ball in place using a magnet. Control animals could see the immobile ball behind a transparent window, but not touch it **(Fig. 2b, left-top)(Video 6)**, whereas experimental animals could also touch the ball and therefore potentially learn its affordance of immobility through direct physical interactions with it **(Fig. 2b, left-bottom)(Video 5)**. After one hour we then removed the window and magnetic tether, allowing both groups to interact with the now mobile ball **(Fig. 2b, right)**. If experimental animals had learned from experience that the ball could not be pushed, one would expect them to show delayed pushing or even a complete disengagement with the ball compared with control animals.

**Fig. 2:**
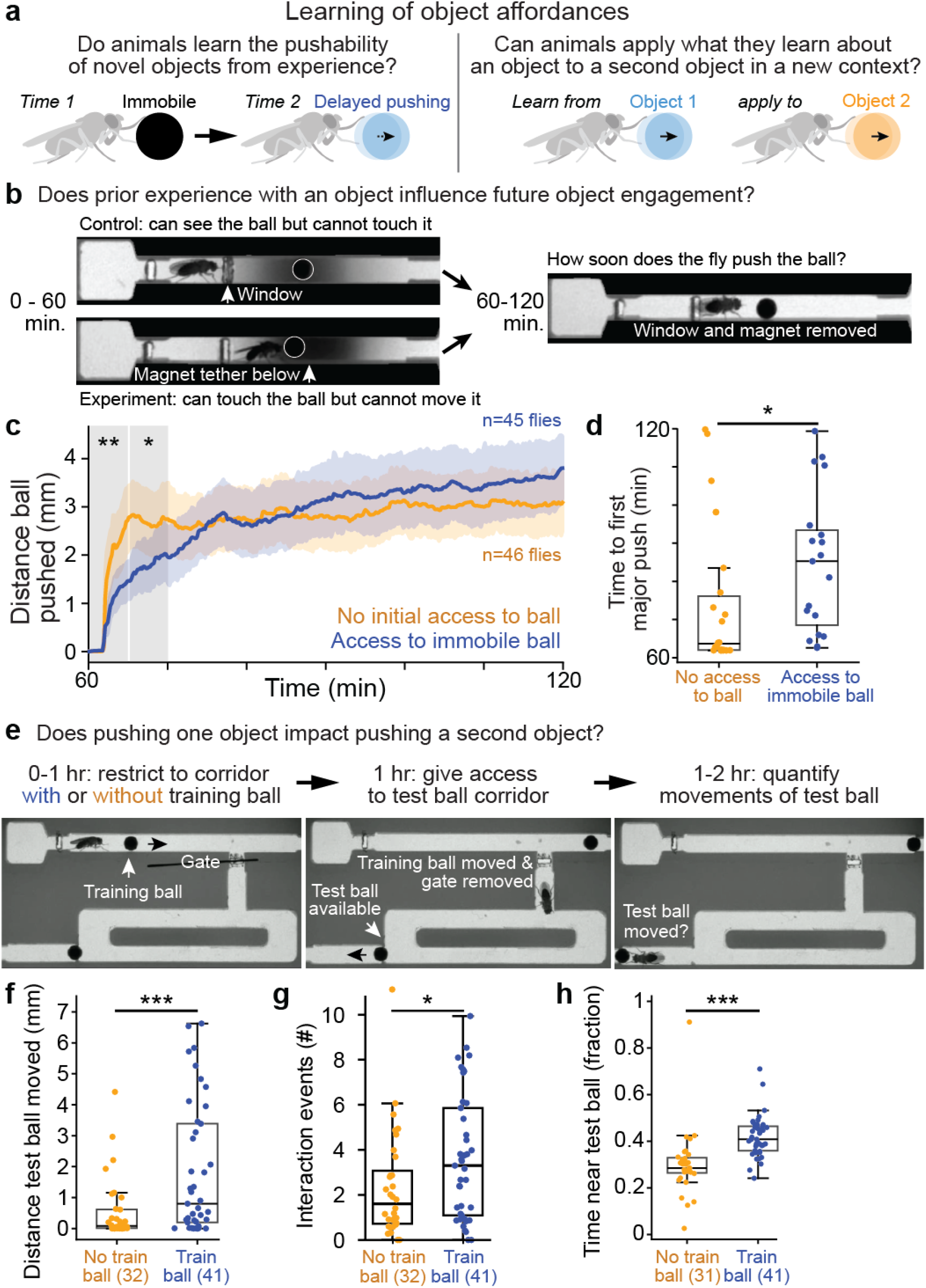
Flies learn the pushability affordance of objects. **(a)** Two experimental paradigms to test the ability of flies to learn object affordances. (Left) If the object is first immobile but then untethered, are flies delayed in learning that it can be pushed. (Right) If flies learn to push one object, are they then able to apply this knowledge to another object in a different context. **(b)** Experimental paradigm to test if flies learn the immobility affordance of balls. For the first hour of the experiment, (top-left) control animals can see the ball but not reach it due to a transparent gate whereas (bottom-left) experimental animals can interact with a ball that is tethered in place by a hidden magnet. Then, after one hour, the window and magnets are removed, allowing both groups of flies to interact with the now mobile ball over the next hour. **(c)** Distance that the ball has been pushed relative to the start position. Shown are mean (solid lines) and bootstrapped 95% CI (shaded regions) for experimental (blue, n = 45) and control (orange, n = 46) animals. Data start at 60 min. when both groups were given access to now mobile balls. Groups were compared using permutation tests at 12 equally-spaced time points with FDR correction (*α* = 0.05); experimental animals pushed significantly shorter distances during the first 10 minutes (0–5 min: *p* = 0.0021; 5–10 min: *p* = 0.032). **(d)** Time until first major ball push (more than 1.2 mm) for control (orange dots, n = 22) and experimental (blue dots, n = 19) animals. Only flies with at least one major push were included. Groups were compared using a permutation test (*p* = 0.0089). Boxplots show the median (center line), interquartile range (IQR; box edges), 1.5 × IQR (whiskers), and individual data points (dots). **(e)** Experimental paradigm to test if affordances of one object can be applied to a second object. (Left) Experimental animals are given a training ball that they can learn to push down a linear corridor. Control animals have no training ball to push. (Middle) After one hour, a gate is removed and the training ball is moved to the corridor end if needed. This gives both experimental and control animals access to a new rectangular corridor with a test ball in one corner. (Right) We quantify animals’ interactions with the test ball over the second hour. **(f-h)** We quantify **(f)** the distance the test ball was moved during interaction events (*p* = 0.0008), **(g)** the number of interactions with the test ball normalized by time spent in the second corridor (*p* = 0.04), and **(h)** the proportion of time flies spend near the test ball after they enter the second corridor (*p <* 0.0001). For control animals n = 31–32 and for experimental animals n = 41. Groups were compared using permutation tests. Boxplots show the median (center line), IQR (box edges), 1.5 × IQR (whiskers), and individual data points (dots). Significance levels: ns, *p* ≥ 0.05; *, *p <* 0.05; **, *p <* 0.01; ***, *p <* 0.001.

Indeed, we found that experimental flies that had been able to interact with the immobile ball took significantly longer to push the newly mobile ball. They pushed shorter distances during the first 10 minutes **(Fig. 2c)** and also exhibited a significant delay until their first major push **(Fig. 2d)**. Importantly, faster pushing in control animals was not due to being startled when the window was removed: walking speeds were not significantly different across experimental and control groups after the window was removed **(Extended Data Fig. 2a-c)**. These results suggest that flies do not solely habituate to the ball, interacting with it more after spending time with it. Instead, flies seem to learn the non-pushability affordance of the ball, and push it less after experiencing it as immobile.

These observations were made in the context in which the affordances of a single object change over time. To more explicitly examine to what extent affordances can be assigned to a particular object, we next asked if flies can apply what they learn about the pushability of one object to a second object in a different context **(Fig. 2a, right)**. Specifically, we designed an experiment in which flies first interacted with a ‘training’ ball in a linear corridor **(Fig. 2e, left)**. After one hour a gate was opened and, if it was not already at the end of the corridor, the training ball was manually pushed clear of the corridor exit. This allowed flies to enter a rectangular track with a ‘test’ ball located in a distant corner **(Fig. 2e, middle)(Video 7)**. Importantly, flies could freely explore the rectangular track without accidentally pushing the ball. Control animals were subjected to the same procedure, but had no initial training ball in the linear corridor. We quantified how each fly pushed the test ball, beginning from the moment it first entered the rectangular track **(Fig. 2e, right)**. We reasoned that, if experimental flies’ interactions with the training ball allowed them to learn the object’s pushability affordance, they would more swiftly and effectively push the test ball compared with control animals. Indeed, we observed that experimental animals pushed the test ball significantly farther than control animals **(Fig. 2f)**, interacted with the test ball more **(Fig. 2g)**, and spent more time near the test ball **(Fig. 2h)**. This shows that flies can generalize the learned pushability affordance of one ball to another.

Taken together, these results suggest that improvements in ball pushing are predominantly due to affordance learning. Motor learning does not play a significant role because flies can perform successful ball pushes in their first expression of a particular movement. Furthermore, unsuccessful pushing of an immobile ball inhibits future pushing attempts, while successful pushing of one ball leads to more efficient pushing of another ball in a different context—both hallmarks of affordance learning.

### A high-throughput neural silencing screen for object manipulation

From our data, it appears that ball pushing dynamics are shaped by at least two phenomena with potentially distinct neural underpinnings. First, an initial retreat that is likely mediated by vision: animals in the dark push with a shorter delay **(Extended Data Fig. 3)**. Second, learning of the ball’s pushability affordance. To gain insights into the neural circuit mechanisms underlying these two processes, we designed a large-scale neural silencing screen. We built a high-throughput experimental system allowing simultaneous recording of 54 corridors **(Fig. 3a)**. With this system, we quantified ball pushing in animals taken from 212 split-Gal4 and Gal4 transgenic driver lines **(Supporting Information Table 1)** for which we constitutively silenced synaptic transmission via expression of tetanus toxin (TNT). We silenced olfactory sensory neurons, mushroom body (MB) intrinsic and extrinsic (e.g., neuromodulatory) neurons implicated in associative learning, neuropeptidergic neurons, central complex neurons involved in navigation, lateral horn neurons involved in innate attractive and aversive behaviors, visual projection neurons, and other neurons including proprioceptors **(Fig. 3b)**. Broadly speaking, we could envision several potential outcomes from neural silencing. First, if multiple circuits perform redundant functions during ball pushing, sparse neural silencing may not have a detectable effect. Second, some circuits may affect behavioral activity levels more globally, driving more or less vigorous movement and interactions with the ball. Third, specific circuits might uniquely regulate different aspects of ball manipulation and affordance learning.

**Fig. 3:**
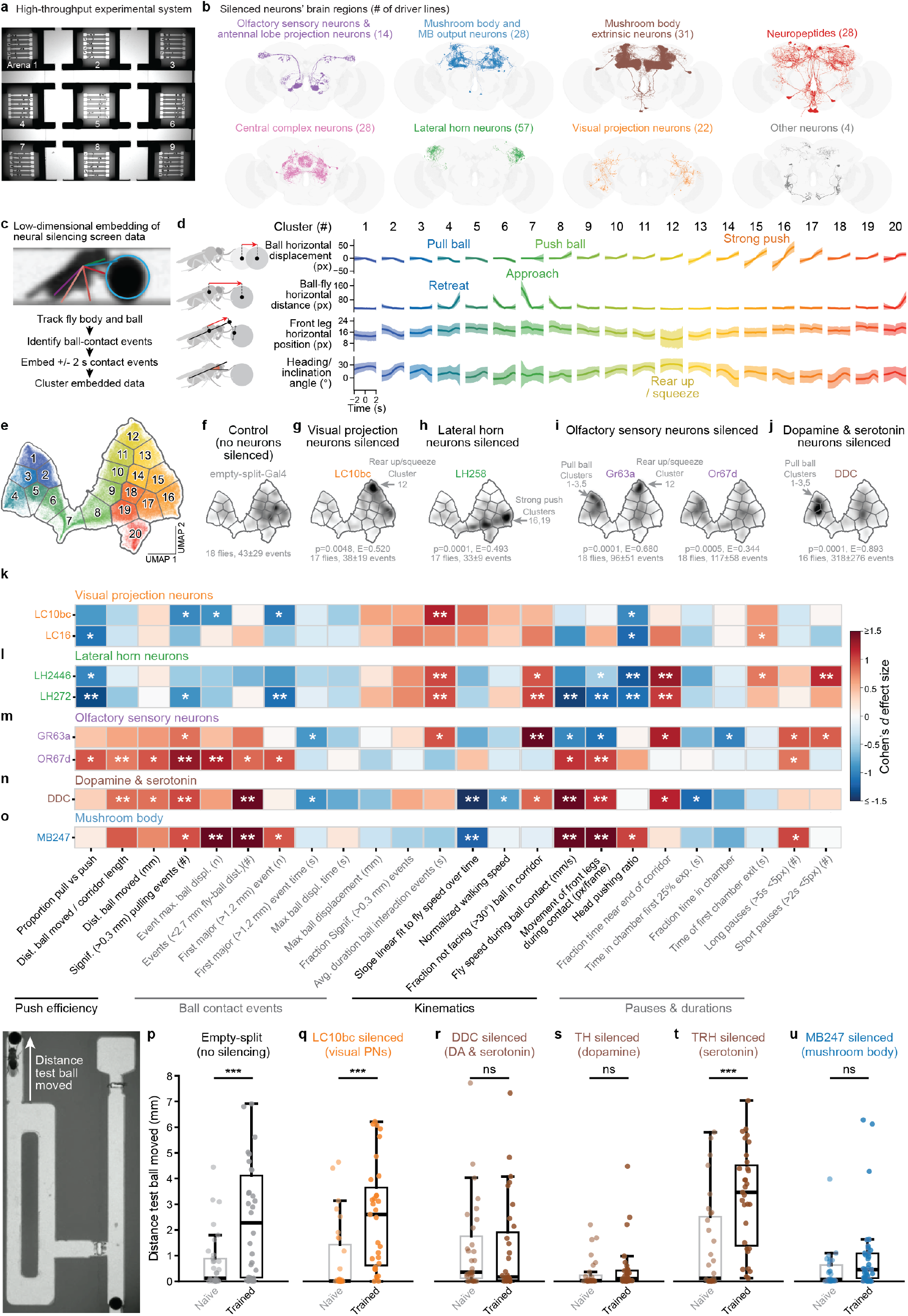
A neural silencing screen reveals roles for visual, olfactory, dopaminergic, and the mushroom body neurons in object manipulation and affordance learning. **(a)** A high-throughput experimental system enables parallel testing of ball manipulation across 54 flies. The system consists of a 3 × 3 array of behavioral arenas, each with 6 linear corridors connected to chambers with custom removable gates. **(b)** Brain regions targeted for neural silencing screen. Shown are connectome (FlyWire) renderings of a subset of targeted neurons (color-coded) on a standard brain template (gray). Numbers in parentheses indicate the number of driver lines tested per region for a total of 212 lines (target: n = 18 flies per line). **(c)** Cropped image from a behavioral recording showing a fly manipulating a 1.5 mm ball during a ‘contact event’. Events are automatically detected based on fly-ball proximity (distance ≤ 2.7 mm for interaction events, ≤ 0.8 mm for contact events). Ball position (circle) and fly leg and head keypoints are annotated (colored lines) using SLEAP. **(d)** Kinematic features are summarized using four metrics: ball displacement following contact onset, fly-ball horizontal distance, front leg horizontal position, and head-body-leg inclination angle. For embedding, tracking data from contact events are temporally-centered and normalized around ball contact (± 2 s time window). Events are embedded using UMAP and clustered into 20 groups (color-coded) based on high-density regions. Highlighted are annotations of fly and ball movements for some clusters. **(e)** UMAP representation of all fly-ball contact events in the silencing screen (n = 197,188 events), color-coded by cluster as in panel d. Left-most clusters (1–7) are associated with weak pushes and pulling whereas right-most clusters are associated with larger pushes. **(f-j)** Enriched (dark) areas in UMAP space for the behaviors of (f) control animals, and those with silenced (g) LC10bc visual projection neurons, (h) LH258 lateral horn neurons, (i) Gr63a- and Or67d-expressing neurons, or (j) dopaminergic and serotonergic neurons. Indicated (gray text) for each driver line are the number of animals tested, the significance of comparison with empty-split-Gal4 control animals, and particularly enriched clusters. **(k-o)** The difference from controls in metrics values for highlighted silenced driver lines targeting **(k)** visual projection neurons, **(l)** lateral horn neurons, **(m)** olfactory sensory neurons, **(n)** neuromodulatory neurons, and **(o)** mushroom body neurons. Columns show manually-sorted behavioral metrics. Cell colors represent Cohen’s *d* effect sizes relative to matched controls (empty-Gal4*>*TNT, empty-split-Gal4*>*TNT, or PR*>*TNT for wild-type), ranging from lower (dark blue; *d* ≤ −1.5) to higher (dark red; *d* ≥ 1.5) values. Significance from permutation tests (FDR-corrected, *α* = 0.05) is indicated as: ns, p ≥ 0.05; *, p *<* 0.05; **, p *<* 0.01; ***, p *<* 0.001. Full statistical results are provided in Supplementary Data. **(p-u)** The distance the test ball was moved for naive (no training ball, light boxes) and pretrained (training ball, dark boxes) animals silencing **(p)** empty-split-Gal4 (control), **(q)** LC10bc-Gal4 (visual projection neurons), **(r)** DDC-Gal4 (dopaminergic and serotonergic neurons), **(s)** TH-Gal4 (dopaminergic neurons), **(t)** TRH-Gal4 (serotonergic neurons), and **(u)** MB247-Gal4 (mushroom body neurons). Naive vs. trained comparisons within each genotype were performed as independent permutation tests with FDR correction (n = 28–33): control *p* = 0.0003; LC10bc *p* = 0.0003; DDC *p* = 0.97; TH *p* = 0.5; TRH *p* = 0.0004; MB247 *p* = 0.08. Boxplots show the median (center line), interquartile range (IQR; box edges), 1.5 × IQR (whiskers), and individual data points (dots). Significance levels: ns, *p* ≥ 0.05; *, *p <* 0.05; **, *p <* 0.01; ***, *p <* 0.001.

We analyzed our neural silencing screen data using two complementary approaches. First, we performed an unbiased exploratory analysis by embedding tracked leg kinematics and ball movements during fly-ball contact events into a low-dimensional UMAP space **(Fig. 3c)** which we then clustered **(Fig. 3d-e)**. Visual inspection of cluster videos revealed distinct aspects of ball manipulation. For example, ball pulling and retreats (like those observed during initial ball interactions) were mainly contained in clusters 1–6 **(Video 8)**. Notably, our nomenclature of ‘ball pulling’ is distinct from that used previously in which flies jump off of larger balls causing them to roll ^21^. Clusters 13– 17 predominantly contained effective ball pushing **(Video 9)**. Control animals with no neurons silenced (i.e., empty-split-Gal4 and empty-Gal4) **(Extended Data Fig. 4a)** showed, as expected, behaviors similar to wild-type animals: they initially retreated and inefficiently pulled the ball but eventually pushed effectively **(Video 10)**. This behavioral diversity is reflected in the enrichment in UMAP space of both the ‘pulling’ clusters 2, 3 and 5 as well as the ‘pushing’ clusters 14, 15, 17 and 18 **(Fig. 3f)**. Across each silenced brain region we observed a diversity of significantly different enrichment patterns in this UMAP space **(Extended Data Fig. 5)(Supporting Information File 2)**, a subset of which we explore below.

In a second analysis, to capture longer-timescale phenomena that might not be apparent during single fly-ball interactions, we designed 31 interpretable metrics related to overall pushing efficiency, the timing of ball contact events, fly kinematics, spatial and temporal tendencies **(Supporting Information File 3)**. To account for covarying factors such as overall activity and speed, we applied dimensionality reduction (PCA and Sparse PCA) to 27 metrics with *<*5% missing values across genotypes, after excluding metrics that were strongly correlated with one another (see Methods). To ensure our results were not sensitive to the specific approach used, we applied 20 different dimensionality reduction configurations to the metrics data, each configuration having been selected via hyperparameter optimization.

We then identified driver lines for which neural silencing resulted in phenotypes that significantly deviated from control animals. From all results **(Supporting Information File 4)**, we retained as robust hits driver lines whose silencing produced metric values significantly different from controls in *>*80% of these configurations (≥ 16/20) **(Supporting Information File 5)**. Hierarchical clustering of the 24 continuous metrics (see Methods) revealed three main groups **(Extended Data Fig. 6)**. Group 1 was characterized by earlier pushing; group 2 showed active but inefficient object interactions often resulting in ball pulling; and group 3 exhibiting more heterogeneous behaviors. In the next sections we focus on hits from the first two groups.

### Silencing visual projection and lateral horn neurons accelerates ball pushing

Two broad lines of evidence suggest that vision plays a key role in driving initial retreat from the object. First, placing wild-type animals in the dark accelerated ball pushing and reduced the latency to the first major push **(Extended Data Fig. 3)**. Second, reducing the reflectivity of the ball’s surface by rusting with ethanol, or through abrasion **(Extended Data Fig. 7a)** also reduced the latency to the first major ball manipulation events **(Extended Data Fig. 7b-c)**. We therefore explored which visual neurons might mediate initial retreat reactions to the ball.

Among the first group of hits from our neural silencing screen—those with earlier ball pushing—we found several lobula columnar (LC) projection neurons known to encode higher-order visual features ^24,25^. Specifically, silencing LC10bc **(Extended Data Fig. 4b)** or LC16 **(Extended Data Fig. 4c)** suppressed initial retreat, resulting in reduced latency to engage in persistent ball pushing. For LC10bc this was quantitatively reflected as a significant enrichment in rearing/squeezing behaviors (cluster 12 in UMAP space)**(Fig. 3g)**. LC10bc-silenced animals also had earlier first major pushing events **(Fig. 3k)**. Similarly, animals with silenced LC16 projection neurons—which are known to be responsible for visually-evoked retreat in response to looming ^26^—exhibited significantly less pulling **(Fig. 3k)**. High-resolution videography confirmed that LC10bc- and LC16-silenced animals begin pushing sooner, often just after the gate is opened and, importantly, were not trivially unable to walk backwards through the corridor **(Video 11)**. Because silencing LC10bc and LC16 yields more immediate engagement with the ball, these neurons may normally mediate visually-driven retreat during initial interactions with the ball. Although the specific function of LC10bc remains to be established, this is consistent with other LC10 neurons encoding visual motion ^27^ and driving retreat when activated ^24,28^.

Silencing specific neurons in the lateral horn (LH), a brain region implicated in innate behaviors ^29^, produced phenotypes that are qualitatively similar to those of LC10bc and LC16. Both LC and LH silencing resulted in more and earlier pushing **(Fig. 3l)**, although the exact behaviors differed somewhat **(Fig. 3h)**. Specifically, silencing LH258 (PV2a1/b1 local interneurons ^30,31^) significantly enriched strong-pushing clusters 16 and 19 in the UMAP space **(Fig. 3h)**, and silencing LH AD1d1 (LH2446) input neurons and PLP-PN1 (LH272) output neurons (of otherwise unknown function ^30^) promoted pushing over pulling **(Fig. 3l)**. These data suggest that these LH neurons may also regulate initial retreat reactions to the ball.

### Silencing neurons in the olfactory periphery accentuates ball pulling

The second group of hits from our neural silencing screen **(Extended Data Fig. 6)** had less efficient object manipulation and increased ball pulling: the opposite phenotype of group 1’s visual projection and lateral horn neurons. Surprisingly, this group includes animals with silenced peripheral olfactory sensory neurons including broad driver lines expressing TNT in olfactory co-receptor IR8a neurons, and neuropeptidergic Mip-1M (myoinhibitory peptide) neurons involved in chemosensing ^32,33,34^—as well as in very sparse driver lines like Gr63a-Gal4 **(Extended Data Fig. 4d)**, which labels chemosensory neurons that detect carbon dioxide ^35,36^, and Or67d-Gal4 **(Extended Data Fig. 4e)** which labels neurons detecting cVA, a male pheromone ^37,38^. We first focused on these two relatively sparse driver lines.

We found that animals with silenced Gr63a neurons had behaviors enriched in UMAP clusters 1–3, and 5 for ball pulling **(Fig. 3i)**. Complementarily, our metrics analysis showed that both Gr63a- and Or67d-silenced animals had more ball pulling events **(Fig. 3m)**. To better understand this unexpected finding, we performed high-resolution videography of Gr63a- and Or67d-silenced animals. There we observed that ball pulling was typically associated with retreat in both control (empty-split-Gal4) and experimental animals **(Video 12)**. This suggests that olfactory neuron-silenced animals may have heightened retreat compared with control animals, indirectly resulting in more ball pulling. We next sought to investigate why this might be the case.

Gr63a and Or67d neurons are sensitive to CO_2_ and cVA respectively, but neither of these chemicals are likely to reach behaviorally relevant concentrations in our assay. CO_2_ avoidance in *Drosophila* requires concentrated emissions from groups of stressed conspecifics ^36^, whereas each fly is alone in the corridor in our experiment. Although cVA is transferred from males to females during mating ^37^, mated females actively eject cVA from their reproductive tract within a few hours of copulation ^39^. Our females were transferred to clean vials without males for 24 hours before testing, providing sufficient time for cVA to be eliminated.

Another potential explanation for the effect of olfactory silencing may be that flies perform scent marking: they may deposit pheromone traces on the ball during initial probing contacts which accumulate, causing animals to become acclimated to the novel object. An inability to detect these neutralizing scent markings due to silencing of specific olfactory sensory neurons might therefore result in more sustained retreat. To test this hypothesis, we manipulated the chemical history of the ball by bathing it in ethanol for 24 hours (to remove odor residues), pre-exposing it to female flies for 24 hours (to add conspecific scents), or sequentially performing first the ethanol bath and then exposure to fly odors. For wild-type animals, none of these conditions altered the rate of ball interactions **(Extended Data Fig. 8a)**, the ratio of pulling versus pushing **(Extended Data Fig. 8b)**, the latency until the first major ball manipulation **(Extended Data Fig. 8c)** or the overall time course of ball pushing **(Extended Data Fig. 8d-f)**. Thus, scent marking likely does not play an important role in fly-ball interactions.

An alternative explanation is that silencing important olfactory pathways may accentuate visual reactions. This may occur for several reasons: silencing olfactory pathways may (i) change the global state of the animals to one of more heightened reactivity, or (ii) induce ‘sensory compensation’ in which suppressing one sensory modality enhances another ^40,41^ possibly by influencing multimodal, integrative central circuits in regions like the MB ^42^. In both cases, silencing IR8a, Gr63a, or Or67d would accentuate visual retreat to ball movements and, because flies’ legs are in contact with the ball, indirectly result in ball pulling. Consistent with this, we found that, in the dark without visual cues, IR8a neuron-silenced animals **(Extended Data Fig. 4f)** pushed the ball substantially faster **(Extended Data Fig. 9a)** and also had a markedly reduced amount of pulling versus pushing **(Extended Data Fig. 9b)**. These data reveal that olfactory sensory neuron-silenced animals are capable of effective ball pushing but that, in the light, olfactory system silencing may heighten visually-induced retreat reactions to the ball.

Taken together, these data suggest that initial retreat reactions to novel objects are bidirectionally regulated by the balanced influence of visual and olfactory sensory pathways: visual pathways (LC10bc and LC16) may normally drive retreat in response to ball movements and delay productive engagement, whereas the spontaneous activity of specific olfactory pathways (IR8a, Gr63a, and Or67d) may normally attenuate these retreat reactions.

### Dopaminergic neurons and the mushroom bodies are indispensable for object affordance learning

To perform effective ball pushing, in addition to reducing their initial retreat reactions, flies appear to learn to apply specific motor programs to push this novel object. One potential framework for explaining this latter process is reinforcement learning (RL). Generally speaking, in RL specific states of the animal or world become paired with particular intentions or actions through reward signaling. These rewards may be explicit, as with the acquisition of food, or implicit and inseparable from the actions themselves ^43^. In the context of learned object manipulation, we hypothesize that the ball’s appearance, texture, scent, and environmental context may become paired (via intrinsic reward-based dopaminergic signaling) with the increased probability of recruiting an innate pushing motor program. Consistent with this possibility, we found that animals with silenced neurons expressing dopa decarboxylase (DDC)—an enzyme required for dopamine and serotonin synthesis that is known to regulate learning in flies ^44^—exhibit ball-pulling phenotypes that place them in group 2 of our clustering of metric values **(Extended Data Fig. 6)**. DDC-silenced animals had some of the most pronounced increases in ball pulling as reflected in the enrichment of UMAP clusters 1–3, and 5 **(Fig. 3j)** as well as in behavioral metrics analysis **(Fig. 3n)**. Importantly, these flies were not trivially inactive: they spent less time in the starting chamber in the early stage of the recordings and moved faster during ball interactions **(Fig. 3n)**.

To directly test the possibility that dopaminergic neurons play a key role in learning object affordances, we next investigated how well animals with neurons silenced performed in our affordance learning experiment. Specifically, we examined the effect of experience with a first, training ball on pushing of a second, test ball **(Fig. 2e)**. As a baseline, we first confirmed that control animals with no neurons silenced (empty-split-Gal4*>*TNT) behaved like wild-type flies **(Fig. 2f)**. Indeed, they more effectively pushed the second test ball after exposure to a training ball **(Fig. 3p)**. Given the evidence from silencing visual pathway neurons, one potential explanation could be that pretrained animals become visually habituated to training ball movements and, thus, retreat less from the test ball. This does not appear to be the case: LC10bc-silenced animals, in which visual retreat is already suppressed, still show a significant improvement in test ball pushing with exposure to a training ball **(Fig. 3q)**. Thus, visual habituation alone cannot account for this pre-training effect.

By contrast, DDC-silenced animals failed to improve test ball pushing after exposure to a training ball **(Fig. 3r)**. Because DDC encodes an enzyme required for the biosynthesis of both dopamine and serotonin ^45^ we next asked which neuromodulatory system was responsible for this deficit. We silenced either dopaminergic or serotonergic neurons using the driver lines TH-Gal4 and TRH-Gal4, respectively. We found that silencing dopaminergic **(Fig. 3s)** but not serotonergic **(Fig. 3t)** neurons disrupted the improvement of ball pushing from pretraining.

Having established that dopaminergic neurons are required for affordance learning, we next asked which brain circuits might employ this dopaminergic signaling. In insects, the mushroom bodies (MB) are the most prominent site for dopamine-dependent associative learning ^46,47^, although other brain regions also experience dopamine-gated plasticity ^48^. Notably, the MB have been studied most extensively in the context of explicit associative learning, such as odor-shock pairing ^49^, rather than operant, self-generated learning as investigated here. Dopaminergic neurons (DANs) project to specific MB compartments where they modulate the mapping of olfactory and visual inputs from Kenyon cells (KCs) onto MB output neurons (MBONs) which then contribute to action selection. DANs are further subdivided into those mediating appetitive (PAMs) versus aversive (PPLs) associations. In our screen, among 28 intrinsic MB driver lines tested, we identified significantly different phenotypes in three KC lines (two were excluded for locomotor defects), and three MBON lines **(Extended Data Fig. 6)**. Among these, MBON-*γ*3/MBON-*γ*3*β*^*′*^1 neurons, which are implicated in appetitive memory acquisition as outputs of the PAM-innervated *γ*3 reward compartment ^50,51^, exhibited a phenotype resembling DDC-silencing **(Extended Data Fig. 6)**. From 34 MB extrinsic and neuromodulatory lines tested, significant hits included PAM-07, PPL1-03, PPL1-04, and a line targeting all PPL1 neurons **(Extended Data Fig. 6)**.

Most strikingly, we found a highly overlapping phenotypic profile between DDC-silenced animals and those with broad but specific silencing of the MB via MB247-Gal4 **(Fig. 3o)**. This driver line primarily targets the *γ* and *α/β* lobes ^52^ **(Extended Data Fig. 4g)**. Importantly, MB247-silenced flies were not simply inactive; on the contrary, they had more ball contact events during which they showed significantly increased speed **(Fig. 3o)**. Unlike DDC-silenced flies, however, MB247-silenced animals also had a significantly delayed first major push **(Fig. 3o)**. When we tested these animals in our affordance learning experiment, the results mirrored those of dopaminergic neuron silencing: exposing MB247-silenced animals to a training ball had a marginal, insignificant effect on pushing of the test ball **(Fig. 3u)**.

Taken together, these neural silencing results support a framework in which two mechanistically distinct processes co-regulate the learning of ball manipulation: (i) sensory-driven reactions to novel objects, and (ii) dopamine- and MB-mediated object affordance learning.

## Discussion

Effectively manipulating objects in the world requires knowing their affordances (i.e., action possibilities) ^10^. Here we explored the neural basis for learning object affordances by investigating how flies interact with a novel spherical object. In contrast with ethological behaviors that are typically studied in flies (e.g., male-female courtship and male-male aggression), this experimental paradigm provides an opportunity to study how animals learn the properties of objects *de novo* as sensation, learning, decision-making, and action unfold in parallel ^53^. We speculate that animals are continuously testing whether novel objects in their environments can be manipulated, allowing them to avoid futile efforts. For example, when navigating towards an enticing food odor, animals may actively probe the compliance of obstacles in their path. In line with this, we found that flies appear to learn the immobility of a magnetically tethered ball: they are delayed in attempting to move it even after it becomes mobile.

Because flies can generate effective pushing strategies without any apparent motor learning, we speculate that they may normally deploy these innate motor programs in natural environments to displace debris or to push through narrow, compliant passages. Notably, although we study female flies exclusively, one effective set of ball pushing movements—rearing and tapping with both forelegs—is similar to a behavior used by males to inspect the species-specific pheromonal profile of prospective mates ^54^. This suggests that this motor program may be part of an innate repertoire of behaviors that can be used across multiple contexts. Thus, we hypothesize that learning to effectively push a novel object requires mapping an existing motor behavior to the sensory context of the object in question. This could explain our observation that after pushing a training ball, flies more effectively push a test ball in a different location.

Which sensory modalities guide this state-action remapping? Our experiments in the dark and our neural silencing screen suggest that vision plays an important role in animals’ experience with these novel objects. Silencing specific visual projection neurons (LC10bc and LC16) greatly reduces the retreat behaviors that flies exhibit during early interactions with the ball and results in more immediate ball pushing. This is consistent with the role of LC16 neurons in visual loom-induced retreat ^26^. Although the behavioral and functional properties of LC10bc neurons have yet to be established, a related cell type, LC10ad, has been implicated in detecting small moving objects ^28^. Therefore, although LC neurons are expected to act as a population rather than as ‘labeled lines’ with each regulating a particular behavior ^24,25^, these particular LC subtypes may play a relatively important role in retreat during initial ball interactions. Further downstream in the central brain, silencing specific lateral horn (LH) neurons—AD1d1 (LH2446) and PLP-PN1 (LH272)—also causes earlier fly-ball engagement, suggesting they may also be part of this sensorimotor retreat pathway. By contrast, we find that silencing specific peripheral olfactory pathways (those expressing the olfactory co-receptor IR8a, or Gr63a and Or67d) increases the degree to which flies pull the ball backwards. We speculate that this is an indirect result of heightened visually-mediated retreat during object interactions. Beyond vision and olfaction, the legs host myriad mechanosensory ^55^, gustatory ^56^, and thermosensory ^57^ neurons which likely also contribute to object sensing. Defining the roles of these other sensory modalities and multi-modal sensory integration in object detection and affordance learning is an important direction for future work.

Our observation that dopaminergic neurons and the MB are critical for learned object manipulation suggests that reinforcement learning may provide a useful mechanistic framework for object affordance learning. This is supported by studies in mammals showing that phasic dopamine can drive the discovery of novel actions ^58^, shape behavioral patterns in the absence of explicit reward ^59^, and act as a gain signal for causal learning between events ^60,61^. In our experiment specific actions like rearing and foreleg tapping precede and enable successful ball pushing and forward locomotion. We speculate that this locomotion may act as an intrinsic reward, driving dopamine signaling which encourages the selection of the same action in future encounters with a ball. Indeed, brain-wide activity of dopamine neurons has been observed during walking ^62^ and turning, where it regulates plasticity in the fly’s navigation system ^48^. Complementarily, dopaminergic neurons associated with aversive learning become active during retreat ^63^. This is consistent with our identification of PPL1 neurons as hits in our neural silencing screen. It also suggests that ball pulling and associated retreat may drive aversive dopaminergic signaling that reduces engagement with the ball—a prediction in line with our simulations **(Fig. 1f)**. Studies in rodents have shown a similar role of striatal dopamine in the degree of engagement with novel objects ^64^. Further work will be required to identify the specific dopamine neuron subtypes that drive affordance learning in flies, and their activity dynamics during this process.

Beyond a mechanistic understanding of affordance learning, the larger question remains of why flies push balls, and do so persistently across multiple trials. What kind of reward encourages this behavior? Rewards may be extrinsic, available in the external world and satisfying a core physiological need (e.g., food, water, and sex). But we do not provide an extrinsic reward in our experiments, suggesting that these behaviors are intrinsically motivated ^43^. The opportunity to explore has been proposed to be intrinsically rewarding ^65^ and this may be achieved through forward displacement of the ball. Such a motivation can also be intuitively linked to the animal’s motivation to escape the arena. Alternatively, curiosity and information seeking have also been proposed to be a ‘manipulatory motive’ even if it does not yield a tangible reward ^66,67^. Curiosity, broadly defined, is an intrinsic motivation to seek information and explore one’s environment ^67,68^. This is closely related to the concept of ‘play’. Some efforts have been made to explore the possibility that insects, including flies ^69^ and bees ^70^, play. However, affirming play behavior requires satisfying formal criteria ^71^, including voluntary engagement in the absence of a competing motivational state. These are conditions that our experimental paradigms do not meet: flies are confined to a narrow corridor and tested when starved and thirsty. Thus, in future work one might explore the concept that flies ‘play’ by asking if fed animals in an open arena repeatedly engage with balls in the absence of an external reward. Irrespective of their underlying motivations, our results establish the fly as a powerful new experimental model for uncovering the mechanistic basis for manipulating objects and learning their affordances—object-action mappings that support navigation and survival in a complex and cluttered natural world.

## Methods

### Fly stocks, husbandry, and experimental preparation

All experiments were performed on female adult *Drosophila melanogaster*, 2–7 days post-eclosion. Animals were raised at 25 °C and 50% humidity on a 12-hour light-dark cycle. Experiments were carried out during flies’ natural activity peaks ^72^.

#### Preparation of flies for experiments

Unless otherwise stated, wild-type animals were starved overnight without access to food or water. Transgenic animals carrying the UAS-TNT transgene exhibited a stronger starvation-induced decline in activity compared with wild-type flies. Therefore, shorter starvation times (4–5 hours) were used for animals in the neural silencing screen. Starvation enhanced the overall displacement of balls relative to fed animals **(Extended Data Fig. 10a-b)**, but did not qualitatively alter the nature of interactions: starved animals showed longer ball-contact interactions and a higher proportion of these contacts resulted in ball-movements **(Extended Data Fig. 10c)**, whereas the number of interactions, their kinematic profiles (i.e., the patterns of leg movements and degree of ball pulling), and the latency until a first major push were comparable across nutritional states **(Extended Data Fig. 10d)**. Starvation therefore modulates the vigor, but not the manner, of ball-pushing. Experiments were performed in the morning following overnight starvation for wild-type animals and in the afternoon otherwise.

#### Experiments with wild-type animals

Wild-type flies were of the PR (Phinney Ridge, gift of M. Dickinson, Caltech USA) strain. Female flies were collected at least 2 days prior to experimentation. Animals were transferred to an empty vial for starvation 22–26 h before experiments. Flies were then tested over 2 h in their morning activity peak around 1–3 zeitgeber time (ZT).

#### Experiments with animals in the neural silencing screen

The neural silencing screen was carried out by crossing UAS-TNT (BDSC#28837) virgin females with Gal4 driver lines selected from 212 Gal4, split-Gal4, mutant, and control lines (Canton-S and Phinney Ridge wild-type strains, empty-Gal4 (BDSC#68384) and empty-split-Gal4 (BDSC#79603)), all crossed with UAS-TNT for consistency. Up to 18 flies were tested for each genotype. All genotypes and cell types are listed in **Table 1**.

### Ball manipulation experimental systems

#### Ball types and ball fabrication

Stainless steel balls (1.5 mm in diameter; McMaster-Carr, 9292K42) were used in most experiments unless stated otherwise. These weigh 13 mg compared with *<*1 mg for a female fly ^73^. In initial experiments, we also tested 1 mm (McMaster-Carr, 9292K29) and 2 mm (McMaster-Carr, 9292K31) diameter balls, weighing 4 mg and 33 mg, respectively.

To test the influence of visual features of the ball on pushing behavior, we altered the standard ball to reduce its reflectivity in two ways. In one case, we used sandpaper to grind down the surface. In the second case, we soaked balls in 90% ethanol for 24 h, altering both the surface and color of the balls through rusting. In both cases, the weights of the balls were not substantially altered.

To test the potential effects of odors on ball manipulation, we altered the standard ball in four ways. First, as a control, we used balls that were never used for experiments before. Second, we briefly washed balls with 70% ethanol to remove odors associated with manufacture and shipping. Third, to add a fly-specific odor, we exposed balls to a group of female flies for 24 h in a petri dish. Fourth, we combined both treatments by first briefly washing the balls in ethanol and then exposing them to female flies for 24 h.

#### Laser cutting experimental arenas

All arenas were designed as .dxf models generated using AutoDesk Fusion 360 (v2606.1.33; Autodesk, Inc., San Francisco, CA, USA). These designs were then laser-cut (TROTEC speedy 400, Austria). Arenas were composed of three layers: a transparent base and ceiling in acrylic (1.57 mm thickness, McMaster-Carr, 8589K12) and a middle section in opaque Polyoxymethylene (POM, 1.6 mm thickness, APSOparts, Switzerland, KT - 02 3200 0104). At a width of 1.6 mm, corridors allowed free movement of the ball while making it impossible for the fly to go around it. Layers were assembled on custom 3D printed holders and held together using rubber stoppers.

Corridors are 17 mm long and book-ended by narrower sections to form a bottleneck that flies, but not balls, can cross. These enter square chambers (4×4 mm) into which flies are initially placed. Visual engravings are added on the side to assist in consistent initial ball positioning, and a removable paper gate is used at the bottleneck to control the start of the experiment.

#### Low-resolution video recording system

The low-resolution recording system is composed of a vertical camera (ImagingSource, Germany, DMK 38UX253) equipped with an infrared filter (Thorlabs, USA, FGL695S). The recorded area is illuminated by an array of infrared LEDs (Solarox, Germany, 58008450500) diffused through a semi-translucent white Polyoxymethylene sheet (POM, 1.6 mm thickness, APSOparts, Switzerland, KT - 02 3200 0104) placed at a distance empirically determined to generate homogeneous rear illumination. This was based on the PiVR design ^74^. Arenas are placed in a 3 × 3 array on a custom grid that also serves as an extra spacer to reduce overheating of arenas from LED illumination. The recording stage is mounted on adjustable posts to ensure that the surface is perfectly flat (checked with a level). The entire system is placed within an opaque enclosure equipped with switchable white light LEDs to either permit or prevent flies from seeing the ball. The camera was operated using custom software. Images are recorded at 29 frames-per-second (fps) using an Arduino-based hardware trigger. Each individual arena location is then cropped and written into a video file. A 3D model of the behavior recording system can be accessed here: https://a360.co/3NUOhEG

#### High-resolution video recording system

Behavioral recordings were acquired at higher resolution using two high-speed cameras (FLIR, Canada, GS3-U3-41C6NIR-C) (one above and one below the arena), each equipped with a macro zoom lens (Computar, Japan, MLM3X-MP). The cameras record the six-corridor arena (including the gating mechanism described above) at 80 fps with a resolution of 2048 × 2048 pixels and an exposure time of 2 ms for a duration of one hour. Infrared illumination is provided above and below the arena using two 850 nm infrared LED ring lights (LDR2-74IR2-850-LA, CCS Inc., Japan), each driven at 24 VDC (6.9 W, 0.29 A) via a digital control unit (PD3-3024-3-EI(A), CCS Inc., Japan). To limit thermal output while maintaining peak illumination intensity during image acquisition, the lights are strobed at a 20% duty cycle (2.5 ms on, 10 ms off; 80 Hz) synchronized to the camera exposure using a custom Arduino script. Uniform visible illumination is achieved using white LED strips (Sloan, Switzerland, FP1-W01D-500) positioned around the arena at a 30° elevation angle on all four sides. Using this high-resolution system, for manual analysis of ball pushing types **(Extended Data Fig. 1)**, we used empty-split-Gal4*>*TNT animals to make results consistent with data from the neural silencing screen.

#### Performing multiple ball manipulation trials for individual animals

To repeat experimental trials for individual animals while minimizing perturbations, we designed a transparent acrylic arm equipped with a small ferrite 3 × 3 mm cylinder magnet (RS Components, UK, 536-1726) as an end effector. The magnet was chosen for its capacity to manipulate a ball in one corridor without influencing those in neighboring corridors. During the recording, we let flies push the ball to the ‘finish’ threshold (10.2 mm from the ball’s start position, allowing for a small margin before the corridor’s end). Upon success, we reposition the ball to its initial position while avoiding contact with the fly. We repeated the ball replacement as many times as possible over 2 hours of recording time. Only animals completing more than four trials are included to enable within-subject comparisons across all four time points.

#### Experimental system to magnetically prevent ball movements

Immobility affordance experiments were performed in the same arenas as regular ball manipulation experiments but consisted of two steps. During the first hour, control animals were kept behind a transparent gate (overhead projector transparency film, ∼ 125 µm polyester sheet) allowing them to see the ball without touching it. Experimental animals were permitted to freely interact with the ball. In both cases, the balls were kept immobile using an horizontal magnet concealed under the back-lighting acrylic plate (Supermagnete, Germany, Q-30-07-2.5-HE). After one hour, the magnets were carefully removed to minimize premature ball displacement, and transparent gates for control animals were removed allowing all flies to freely interact with the now mobile balls for one more hour. If any significant movements of the ball were caused by magnet removal, these data were discarded.

#### Experimental system to examine the influence of a training ball on test ball manipulation

Experiments exploring the generalization of ball pushability affordances were performed in arenas with two compartments. One is similar to the corridor design with a starting chamber and a linear corridor. The second is a rectangular corridor designed to allow animals to freely walk without having to encounter the ball. The test ball is placed at one end of the rectangle, aligned with the corridor wall. Both ends of the rectangle were enlarged to minimize accidental collisions with the ball. The linear corridor and rectangular corridor are bridged by a bottleneck designed to prevent the training ball from entering the rectangular arena.

#### High-throughput neural silencing screen

Neural silencing screen recordings were performed in three batches per day, ranging from one hour before to two hours after their evening peak of activity (between 11 and 13 ZT). Flies were recorded for one hour and genotypes were selected so that each tested line was tested over at least two days, in shuffled locations on the 3 × 3 layout, using flies from at least two distinct crosses to minimize confounding factors.

### Behavioral data analysis

#### Tracking fly and ball positions

To detect fly-ball interactions in low-resolution video data, each fly’s thorax and the center of the ball were tracked using a shallow trained network (MobileNetV2 ^75^) with SLEAP ^23^ version 1.3.0. For ball center tracking, three points at the edge of the ball were labeled for each training frame, and the center was computed using the following formula:

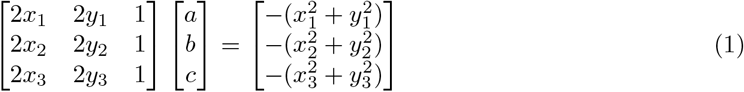

Where (*x*_1_, *y*_1_), (*x*_2_, *y*_2_), and (*x*_3_, *y*_3_) are the coordinates of the three points on the circle. After solving this system for *a, b*, and *c*, the center coordinates (*x*_*c*_, *y*_*c*_) and radius *R* are given by:

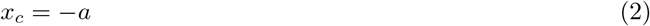

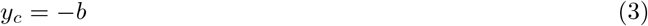

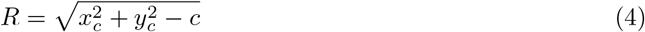

This method ensures an accurate center location even under noisy conditions.

#### Fly body tracking

Fly body 2D pose estimation was performed using SLEAP version 1.3.0 ^76^. We trained a second model using the LEAP backbone ^77^ and labeled keypoints on the head, thorax, abdomen, front legs, middle legs, hind legs and wings. Lateral symmetry was also specified in the training configuration. To improve model performance, raw videos were preprocessed by resizing to a standard size template, applying histogram equalization for brightness discrepancy correction, and cropping the corridors to further reduce ambient noise.

#### Identifying fly-ball interaction events

We computed interaction events as periods in which flies were close to the ball (thorax position ≤ 2.7 mm from ball center) for at least 2 s. If two events were separated by less than 4 s, they were considered one event and grouped together. Events are further sub-classified by ball displacement: significant events indicate ball movements above noise level (≥0.3 mm); major events indicate substantial pushes (≥1.2 mm).

#### UMAP analysis

Contact events were defined as time intervals during which the Euclidean distance between the ball and the tip of the nearest foreleg was *<*13 px. This threshold was determined empirically by visually inspecting video frames and identifying the distance value that best corresponds to apparent physical contact between the tip of the foreleg and the surface of the ball. Consecutive intervals separated by gaps shorter than 0.5 s were merged, and merged events with durations *<*0.5 s were excluded. For each contact event, we extracted a ±60-frame window centered on event onset and computed four time-varying variables: (1) ball displacement along the corridor relative to event onset, (2) ball–fly distance along the corridor, (3) anterior–posterior position of the midpoint between the front leg tips along the body axis, and (4) fly heading (0° = forward). Events in which the ball was displaced by less than 8 px between the start and end of the window were excluded from further analysis.

Each event was represented as a flattened 480-dimensional vector (120 frames × 4 variables) and embedded into two dimensions using UMAP. To eliminate redundant symmetries in the embedding space (e.g., the same type of movement occupying the left and right parts of the UMAP space), we used a modified Euclidean distance metric in UMAP: dist(**x**_*i*_, **x**_*j*_) = min ∥**x**_*i*_ − **x**_*j*_∥_2_, ∥**x**_*i*_ − mirror(**x**_*j*_)∥_2_, where mirror(**x**_*j*_) denotes the mirrored version of event *j* about the corridor midline.

For visualization and to examine how contact events varied across the embedding space, we partitioned the embedded points into 20 clusters using K-means. Cluster indices were ordered by solving a traveling salesman problem to minimize the total distance between consecutive cluster centers.

#### Metric calculations

We computed interaction event-based metrics:

- **Number of interactions**: In cases where the ball access was differentially restrained (i.e., in the affordance generalization experiment) or when the analysis was restricted to the time before reaching the end of the corridor, this number was normalized by the total time available for interactions. Associated metric in the screen: *Events (<2*.*7 mm fly-ball dist*.*)(#)*
- **Number and ratio of significant interactions**: Interactions leading to ball displacement greater than noise (*>*5 px/0.3 mm).
- **Fraction Signif. (***>***0.3 mm) events**: Proportion of significant interactions among all interactions.
- **Major event**: Event at which the ball was displaced by at least 20 px (1.2 mm), a threshold determined empirically from the observed distribution of ball displacements during contact events. Associated metrics in the screen : *First major (>1*.*2 mm) event(#), First major (>1*.*2 mm) event time (s)*
- **Maximum event**: Event at which ball displacement was maximal for a particular individual. Associated metrics in the screen: *Event max. ball displ. (#), Max ball displ. time (s)*.
- **Final event**: Event (if any) at which the ball was brought to the maximum possible distance (170 px/10.2 mm for standard experiments).
- **Signif. (***>***0.3 mm) pulling events (#)**: Number of significant events leading to backward movements of the ball.
- **Proportion pull vs push**: Ratio of the number of significant events resulting in the ball being pulled over the total number of significant events.

*Note*: For all metrics related to particular events in time, both the index of the event and the associated time were computed.

We also computed binary metrics corresponding to the presence or absence of key interactions during the recording. For instance, *has finished* corresponds to whether the fly brought the ball to the end of the corridor, *has significant* indicates whether a ball movement above noise level occurred and *has major* indicates whether a major event occurred.

In addition to the specific interaction-related metrics above, we computed metrics related to activity levels:

- **Overall interaction rate**: Total interaction frequency in events/seconds.
- **Slope linear fit to fly speed over time**: Linear change in fly speed over time, computed as the slope of speed over time.
- **Normalized walking speed**: speed adjusted for available space the fly has to move, computed using the ball’s position in the corridor.
- **Fly speed during ball contact (mm/s)**: Average speed of the fly when it is interacting with the ball.

Ball manipulation efficiency was quantified through:

- **Dist. ball moved**: Total cumulative distance the ball was displaced across all interactions.
- **Dist. ball moved / corridor length**: Ratio of total distance moved to maximum distance achieved, indicating manipulation efficiency (values near 1 suggest direct, efficient movements; higher values indicate repeated back-and-forth manipulations).
- **Max ball displacement (mm)**: Furthest distance the ball reached from its starting position.

Behavioral patterns during interactions were assessed using:

- **Movement of front legs during contact (px/frame)**: Average motion energy of front legs during interaction events, computed as the sum of squared speed differences for leg keypoints. Higher values indicate more energetic, potentially inefficient leg movements.
- **Head pushing ratio**: Proportion of contact frames where the head is closer to the ball than the legs, distinguishing head-based from leg-based manipulation strategies.

Pause and arrest behaviors were characterized across multiple timescales:

- **Short pauses (***>***2s** *<***5px) (#)**: Number of brief pauses, potentially reflecting momentary hesitation.
- **Long pauses (***>***5s** *<***5px) (#)**: Number of pauses, marking behavioral arrest.
- **Total pause duration**: Total time spent pausing. The short pause threshold was used to capture as much pausing as possible.

Spatial strategies and persistence were quantified by:

- **Time of first chamber exit (s)**: Latency to permanently leave the starting chamber, indicating exploration onset.
- **Chamber time and ratio**: Total time and proportion of experiment spent in the starting chamber. Associated metric: *Fraction time in chamber*.
- **Time in chamber first 25% exp. (s)**: Time spent in chamber during the first 25% of the experiment, capturing delay to explore the corridor.
- **Avg. duration ball interaction events (s)**: Average duration of individual interaction events, reflecting how sustained manipulation attempts are.
- **Fraction time near end of corridor**: Fraction of time spent at the corridor end (goal area), indicating goal-directed persistence after potential task completion.
- **Fraction not facing (***>***30**^*°*^**) ball in corridor**: Proportion of time when body orientation deviates *>*30^*°*^ from direct line to ball, potentially indicating distraction, exploration, or pulling-based strategies.

#### Consistency analysis to identify metrics-based significant phenotypes

To identify genotypes with significantly altered ball-pushing behavior, we performed statistical analyses using Principal Component Analysis (PCA) and its sparse variant to reduce the dimensionality of correlated behavioral metrics while preserving biological signal and minimizing technical noise.

##### Data cleanup and correlation analysis

We first excluded metrics with *>*5% missing values, then removed samples with any remaining missing values. We verified that excluded samples were distributed evenly across genotypes, confirming that this procedure did not disproportionately impact any particular driver line. To reduce metric redundancy, we used two parallel metric selection strategies. For correlation-based selection, we computed pairwise Pearson correlations and performed hierarchical clustering (average linkage) on correlation distance matrices (1− |*ρ*|).

From each cluster, we selected the representative metric with the lowest objective score (correlation degree + missingness penalty − bootstrap stability bonus), where correlation degree quantifies how many other metrics exceed |*ρ*| *>* 0.8, and bootstrap stability reflects selection frequency across 100 resampled datasets. For family-based selection, metrics were assigned to behavioral families (timing, intensity, rates/ratios, state/success, persistence/dynamics, orientation/kinematics) based on semantic content, and the same correlation-clustering approach was applied within each family to preserve cross-family diversity while removing within-family redundancy. Of these 27 metrics, 3 binary metrics (has finished, has significant, has major) were excluded from hierarchical clustering and visualization because binary metrics are not suitable for Pearson correlation-based distance measures. The remaining 24 continuous metrics are shown in **Extended Data Fig. 6** and **Fig. 3**.

##### PCA configuration and hyperparameter selection

To explore sensitivity to model configuration and select optimal Sparse PCA hyperparameters, we performed an exhaustive grid search across two metric sets (correlation-based and family-based selection) and two decomposition methods (standard PCA and Sparse PCA), yielding four conditions in total. For standard PCA, the number of components was varied over *n* ∈ {7, 10, 12, 15, 20}. For Sparse PCA, three parameters were searched: number of components (*n* ∈ {7, 10, 12, 15}), the sparsity regularization penalty (*α* ∈ {0.01, 0.05, 0.1, 0.5, 1, 2, 5, 10}), and the ridge penalty (*α*_ridge_∈ {0.01, 0.1, 1.0}). Each parameter combination was scored using a weighted composite of explained variance ratio (PCA: 40%, Sparse PCA: 30%), reconstruction error (20%/15%), component interpretability—fraction of components with 10–90% zero loadings (20% each), sparsity targeting 30–70% zero loadings (10%/25%), and algorithm convergence (10% each). All scores were penalized by a Bayesian Information Criterion (BIC) term (weight 0.15 for PCA, 0.18 for Sparse PCA) to discourage overfitting through excess components. The top 5 parameter sets per condition were retained, yielding 20 configurations in total used for robustness assessment.

##### PCA runs

For each of the 20 configurations (5 best-scoring parameter sets ×4 conditions), data were robust scaled and the appropriate method (PCA or Sparse PCA) applied to the corresponding metric subset. Genotype-control comparisons used tailored controls: Empty-Gal4*>*UAS-TNT for GAL4 lines, empty-split-Gal4*>*UAS-TNT for split-GAL4 lines, and PR (wild-type parental strain)*>*UAS-TNT for mutant strains. For each comparison, we applied three complementary statistical tests: (1) permutation testing (10,000 iterations) comparing multivariate centroids (mean PC score vectors) in reduced component space; (2) Mahalanobis distance measuring variance-corrected separation between centroids while accounting for covariance structure; and (3) univariate Mann– Whitney *U* tests on individual PC scores. All p-values were FDR-corrected (*α* = 0.05). A genotype was considered a hit if the multivariate tests (permutation and Mahalanobis) and at least one individual PC (Mann–Whitney) each yielded significance.

Across the 20 configurations, each genotype received a consistency score (proportion of configurations yielding significance). Genotypes with consistency *>* 80% (16/20 configurations) were classified as robust hits, yielding 29 genotypes prioritized for further analysis.

##### Detailed metrics analysis

For phenotypic characterization, these 29 hits were compared to matched controls on all 24 continuous behavioral metrics using permutation tests (10,000 iterations, FDR-corrected, *α* = 0.05), with effect sizes quantified as Cohen’s *d* (clipped to [−1.5, 1.5]). Genotypes were hierarchically clustered by Euclidean distance in Cohen’s *d* space; metrics were clustered by Pearson correlation distance (1 −| *ρ*|), both using Ward linkage (scipy.cluster.hierarchy). The resulting genotype × metric matrix was cut into three groups (fcluster, criterion maxclust), with within-group ordering preserved from the full dendrogram leaf order. This clustering was performed on raw metric effect sizes, independently of the PCA pipeline described above.

### Modeling ball manipulation experiments

We implemented a discrete-time, one-dimensional simulation to model fly-ball interactions. Each simulated fly occupies integer positions starting at *x*_0_ = 0; time advances in uniform integer steps. Movement is represented as a bounded random walk: at each timestep the fly’s position either increases or decreases, each with probability *p* = 0.5, within the bounds of 0 and the current ball position, *b*_*t*_.

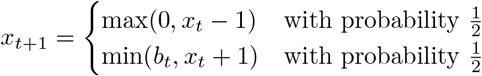

When the fly is at the same position as the ball, a push is attempted which succeeds with probability 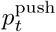. A successful push increases the ball’s position by one. Additionally, a reinforcement mechanism was implemented by which a successful or unsuccessful push would, respectively, increase or decrease the probability of a future successful push, by an amount Δ.

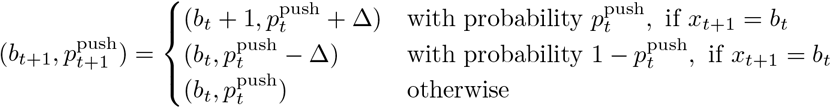

Multiple independent simulations were performed (*N* = 53 in Fig. **1**) and the ball’s trajectory during each simulation was recorded. The Δ parameter was set either to 0 or 0.08 to test the effect of learning on the resulting distributions of ball trajectories. Simulations were implemented in Python in this script.

### Data plotting and statistical analyses

Data were analyzed and plotted using Python (v3.11).

Summary metrics were plotted as boxplots. The middle line represents the median, while the upper and lower limits of the box are the 25 and 75% quantiles. The whiskers are the maximum and minimum values of the data that are, respectively, within 1.5 times the interquartile range over the 75th percentile and under the 25th percentile. Raw data are superimposed as jittered dots.

Statistical significance is indicated by asterisks: \**p <* 0.05; \*\**p <* 0.01; \*\*\**p <* 0.001. Comparisons passing significance threshold are annotated; the absence of asterisks indicates no significant difference. Detailed statistics for each figure and supplementary figure can be found on **Supporting Information File 6**.

### Immunofluorescence imaging of brains and ventral nerve cords

We performed confocal imaging on a subset of driver lines tested in our neural silencing screen by crossing these lines with UAS-Spaghetti Monster GFP (BDSC#62147).

#### Tissue dissection and fixation

Brains and ventral nerve cords (VNCs) were dissected from 2–3 days post-eclosion (dpe) female flies in PBS and fixed for 20 min in 4% paraformaldehyde in PBS at room temperature.

#### Immunostaining

After fixation, tissues were washed 2–3 times in PBS with 1% Triton X-100 (PBST) for 10 min each, then incubated overnight at 4 °C with primary antibodies diluted in PBST supplemented with 5% Normal Goat Serum (PBST-NGS):

- Rabbit anti-GFP (1:500; Thermo Fisher Scientific, RRID: AB 2536526)
- Mouse anti-Bruchpilot/nc82 (1:20; Developmental Studies Hybridoma Bank, RRID: AB 2314866)

Tissues were rinsed 2–3 times in PBST for 10 min each, then incubated for 4 h at room temperature with secondary antibodies diluted in PBST-NGS:

- Goat anti-rabbit conjugated with Alexa Fluor 488 (1:500; Thermo Fisher Scientific)
- Goat anti-mouse conjugated with Alexa Fluor 633 (1:500; Thermo Fisher Scientific)

Finally, tissues were rinsed 2–3 times in PBST for 10 min each and mounted onto slides with bridge coverslips in SlowFade mounting medium (Thermo Fisher Scientific, USA, S36936).

#### Image acquisition

Samples were imaged on an Olympus FV4000 laser scanning confocal microscope (IX83 stand) using a UPLXAPO 10 × objective (0.4 NA, 3.1 mm WD). Images were acquired at 2048 × 2048 pixels (0.22 × 0.22 µm XY pixel size) over 103 Z-slices at 2.84 µm intervals (total stack depth ≈ 290 µm), with a 16-bit dynamic range and galvano scanners in sequential acquisition mode. Channel settings were as follows: CH1 (Alexa Fluor 488): 488 nm excitation at 3.41% laser transmissivity; CH2 (Alexa Fluor 633): 640 nm excitation at 3% laser transmissivity, 50 µm pinhole. No frame averaging was applied (integration count = 4; Kalman: 4 accumulations, average = 6). Laser intensities and PMT gains were kept constant across all driver lines to allow direct comparison of GFP expression levels.

#### Image processing

Maximum intensity Z-projections (MIPs) were computed from the raw multichannel TIFF stacks using a custom Python script (`Zstack renderer.py`) available in this repository. Briefly, each TIFF file was loaded using`tifffile` and reshaped into a canonical (`C, Z, Y, X`) array, accommodating both single- and multi-channel inputs as well as variable axis ordering. The MIP was computed by taking the voxel-wise maximum along the Z-axis, yielding a single (`C, Y, X`) projection per image. For display purposes, pixel intensities were contrast-stretched between user-defined lower and upper percentile bounds, followed by optional gamma correction and brightness scaling, before being clipped to the [0, 1] range. To ensure comparability across lines, all contrast enhancement parameters (*γ*, brightness, and percentile bounds) were held constant throughout the dataset.

Each brain and VNC volume was then registered to the nc82 channel of an empty-Gal4 reference brain (lacking transgene expression), and subsequently warped to the canonical JRC2018 female brain template ^78^ to ensure consistent anatomical alignment across all lines.

### Use of Large Language Models

Large language models (LLMs) were used in the preparation of this manuscript. Specifically, LLMs were used as coding assistants during the development, refactoring, debugging, and optimization of data analysis and tracking pipelines. LLMs were additionally used for writing assistance during drafting and revision of the manuscript. All LLM-assisted outputs were reviewed, verified, and edited by the authors, who take full responsibility for the content of this manuscript.

## Data availability

Data are organised thematically in datasets containing raw .h5 tracking data, .mp4 grid videos and .feather/ .parquet processed datasets. These can be used to reproduce figures and statistical analyses using the code repository below.

Links to the datasets:

- Affordance experiments: https://doi.org/10.7910/DVN/91R87T
- Neural silencing screen: https://doi.org/10.7910/DVN/SPBKKJ
- Wild-type and supplementary experiments: https://doi.org/10.7910/DVN/VB4UI5
- High-resolution videos of fly interactions with the ball: https://doi.org/10.7910/DVN/YU0DPS

## Code availability

Analysis code is available at: https://github.com/NeLy-EPFL/ballpushing_utils

## Extended data figures

**Extended Data Fig. 1:**
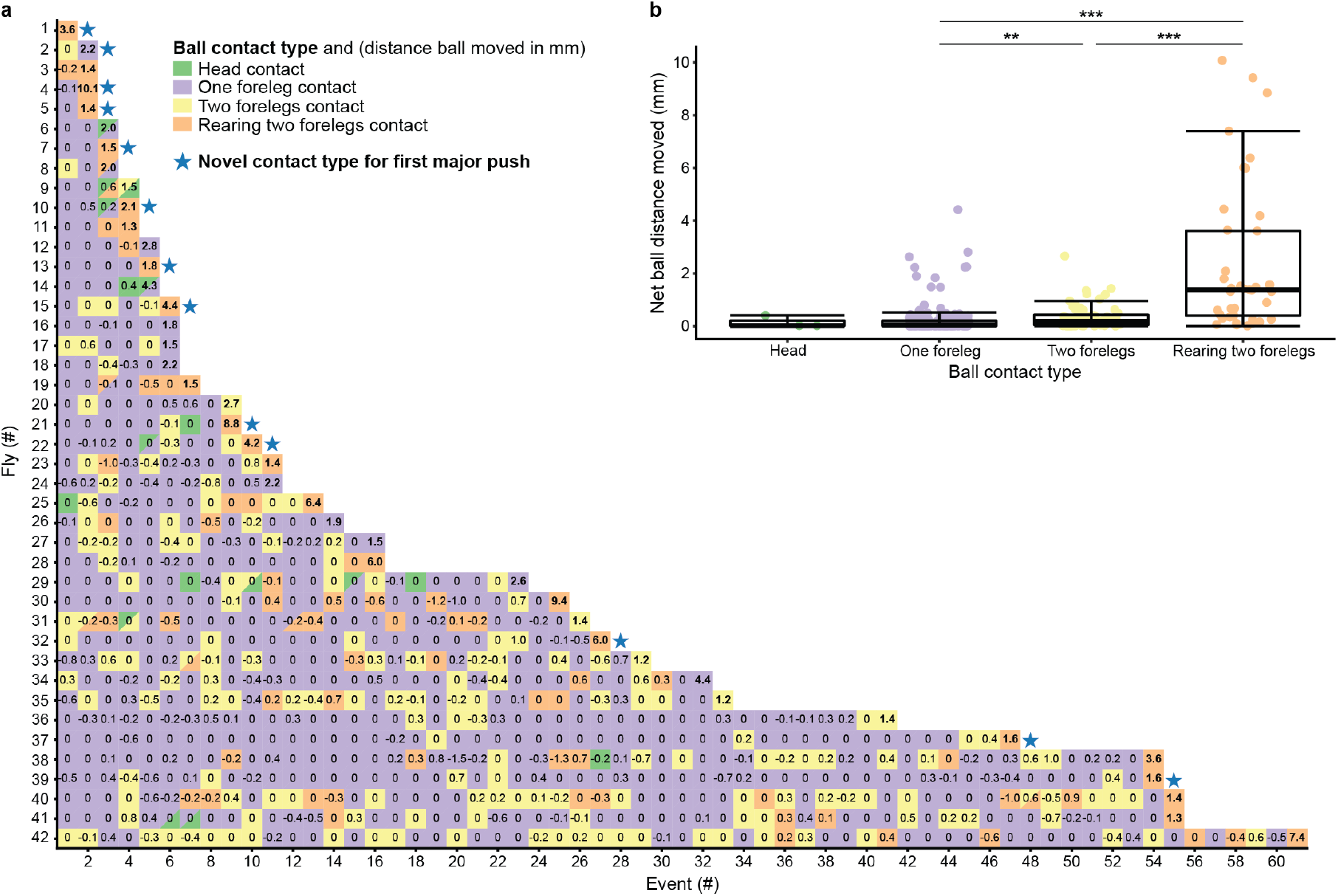
Ball pushing contact types and their pushing efficacy. **(a)** Contact events were classified across event indices for 42 Empty-split-Gal4*>*UAS-TNT animals, until the first major event (ball distance pushed ≥ 1.2 mm) was achieved. Ball contact types are indicated (color-coded). The distance the ball was moved (in mm) is overlaid; bold font indicates a major event. Hybrid events combining two contact types are shown (cells split diagonally in two). Blue stars denote if the first major push is a novel contact type, not previously exhibited by that individual fly. Flies are sorted by the index of the first major event. **(b)** Net ball distance moved during each pushing event grouped by ball contact type. Data are plotted as boxplots with raw data superimposed as jittered scatter plots (n = 3, 150, 68, 40 contact events for ‘head’, ‘one foreleg’, ‘two forelegs’, and ‘rearing two forelegs’, respectively). Rearing pushes caused the ball to move significantly further than other pushing types (permutation test, *p* = 0.0003 vs both one foreleg and two foreleg). Head pushes were not included in statistical comparisons because of the low sample size. Groups were compared using permutation tests (*α* = 0.05). Boxplots show the median (center line), interquartile range (IQR; box edges), 1.5×IQR (whiskers), and individual data points (dots). Significance levels: ns, *p* ≥ 0.05; *, *p <* 0.05; **, *p <* 0.01; ***, *p <* 0.001.

**Extended Data Fig. 2:**
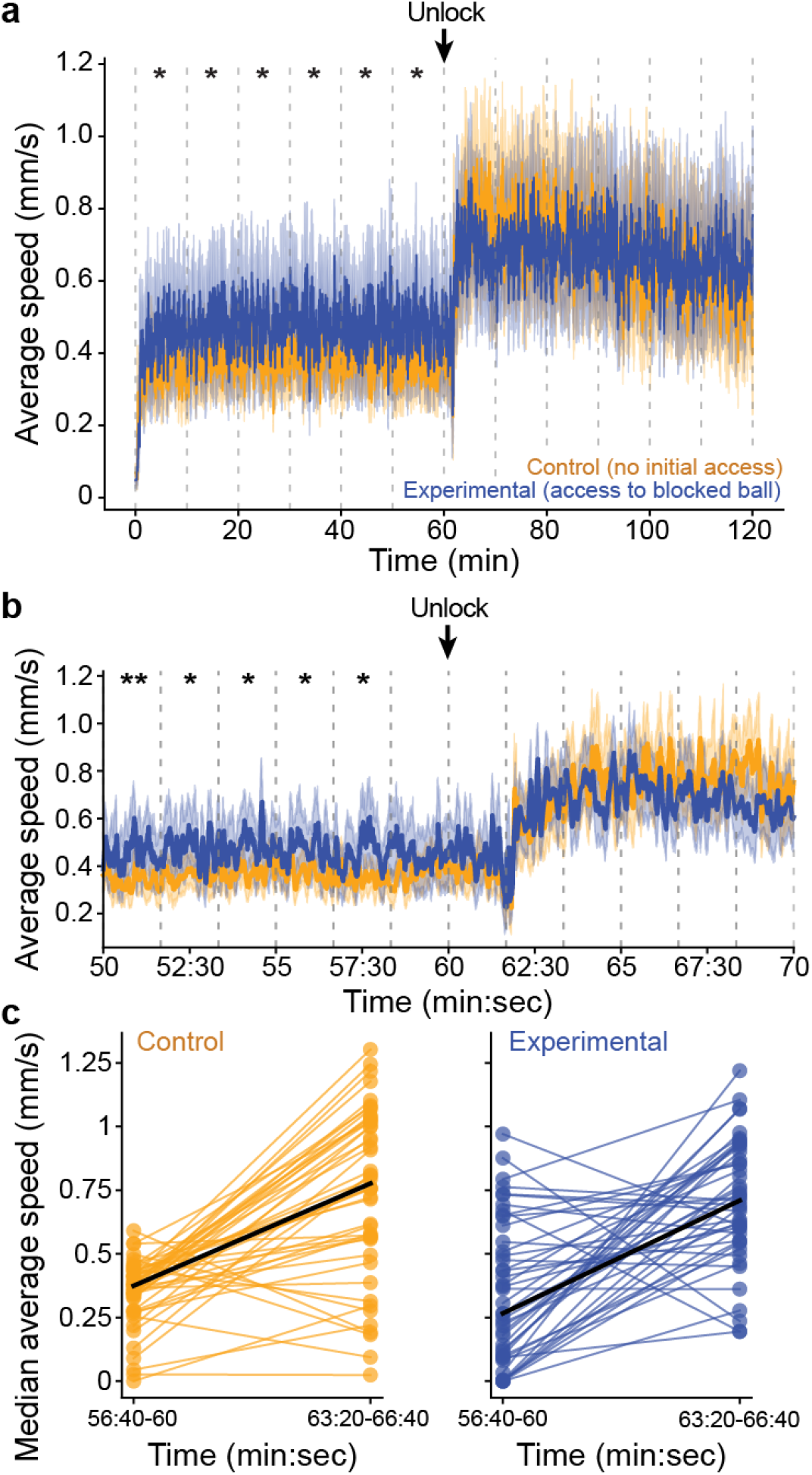
Fly speed and interaction rates during ball immobility affordance experiments. **(a-b)** Average walking speed of control (orange, n = 46) or experimental (blue, n = 45) flies for **(a)** the entire, or **(b)** 20 minutes surrounding the time of gate opening and magnet removal (black arrow, ‘Unlock’). Shown are mean (solid line) and bootstrapped 95% confidence intervals (shading). Data are binned into **(a)** 10 minute, or **(b)** 100 s segments and compared using permutation tests with FDR correction (*α* = 0.05). In **(a)**, experimental animals walked significantly faster than controls in the first six bins (time bins 1–6, all FDR-corrected *p* ≤ 0.019), but not thereafter (bins 7–12, all *p* ≥ 0.48). In **(b)**, experimental animals walked significantly faster than controls in the first five bins prior to removal of the magnet (bins 1–5, all FDR-corrected *p* ≤ 0.021), but not after (bins 6–12, all *p* ≥ 0.051). **(c)** Changes in average walking speed for individual animals following gate opening and magnet removal. Shown are each fly’s median speed in the pre-unlocking (56:40–60:00 min) or post-unlocking time window (63:20–66:40 min) connected by lines. Indicated are the medians for each condition (black lines). The change in walking speed upon gate opening did not differ significantly between groups (two-sample permutation test, *p* = 0.70). All statistical tests used permutation tests with FDR correction (*α* = 0.05). Significance levels: ns, *p*≥ 0.05; *, *p <* 0.05; **, *p <* 0.01; ***, *p <* 0.001.

**Extended Data Fig. 3:**
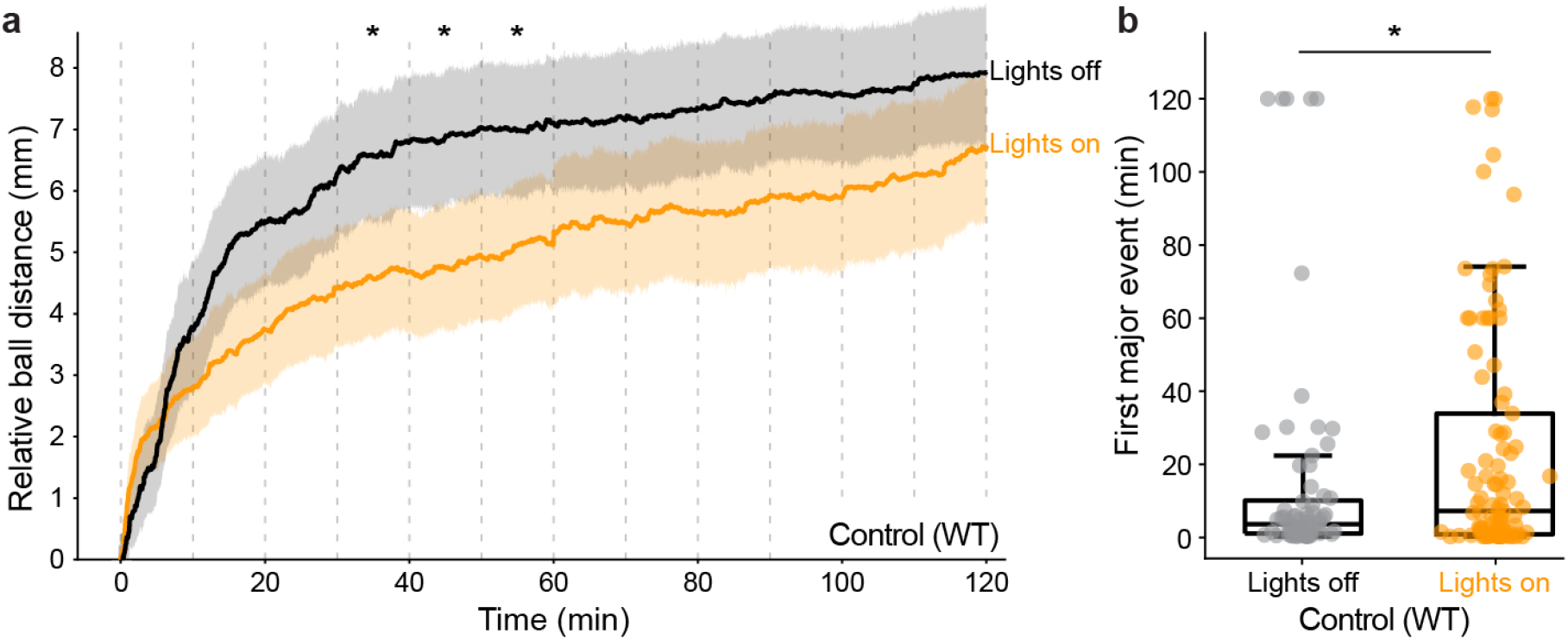
Ball pushing of wild-type animals under different illumination conditions. **(a)** Distance the ball was pushed over 2 h of the experiment relative to the ball’s start position. Shown are data either with (orange) or without (black) illumination plotted as means and bootstrapped 95% confidence intervals. Time-series data were binned into 10 min. segments and compared using permutation tests with FDR correction (*α* = 0.05). Animals in the dark pushed the ball significantly farther in bins 4–6 (30–60 min; all FDR-corrected *p* ≤ 0.047), but not in earlier or later bins (all *p* ≥ 0.057). **(b)** Time until the first major ball interaction event (displacement *>* 1.2 mm) for lights off (gray, n = 72) or on (orange, n = 101 flies). Animals in the dark had a significantly earlier first major push (permutation test, *p* = 0.039). Groups were compared using permutation tests (*α* = 0.05). Boxplots show the median (center line), interquartile range (IQR; box edges), 1.5 × IQR (whiskers), and individual data points (dots). Significance levels: ns, *p* ≥ 0.05; *, *p <* 0.05; **, *p <* 0.01; ***, *p <* 0.001.

**Extended Data Fig. 4:**
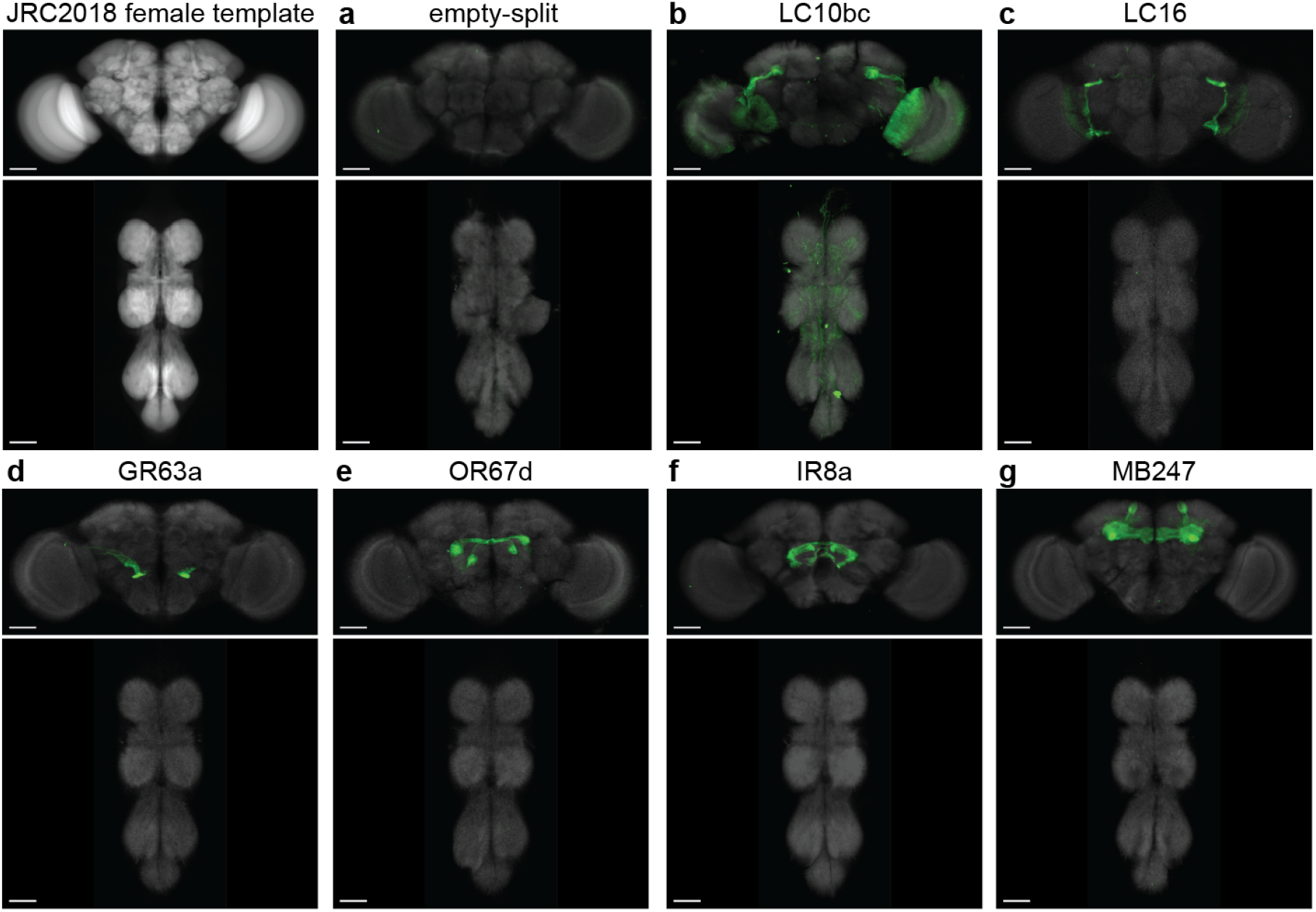
Immunofluorescence-stained confocal images of driver lines highlighted in the neural silencing screen. **(a-g)** Maximum intensity projection (MIP) images of confocal volumes for **(a)** empty-split-Gal4, **(b)** LC10bc-Gal4, **(c)** LC16-Gal4, **(d)** Gr63a-Gal4, **(e)** Or67d-Gal4, **(f)** IR8a-Gal4, and **(g)** MB247-Gal4 registered to the JRC2018 female brain template. The top-left panel shows the reference template (JRC2018; nc82 channel) for context. All remaining panels display MIPs of brains (top rows) and ventral nerve cords (bottom rows) for each Gal4 driver line tested. Green and gray signals, respectively, reflect the expression pattern of UAS-Spaghetti Monster GFP (BDSC#62147) and nc82 labeling of presynaptic active zones, serving as a neuropil reference. All images were acquired and processed using identical parameters (see Methods). Scale bars are 50 µm.

**Extended Data Fig. 5:**
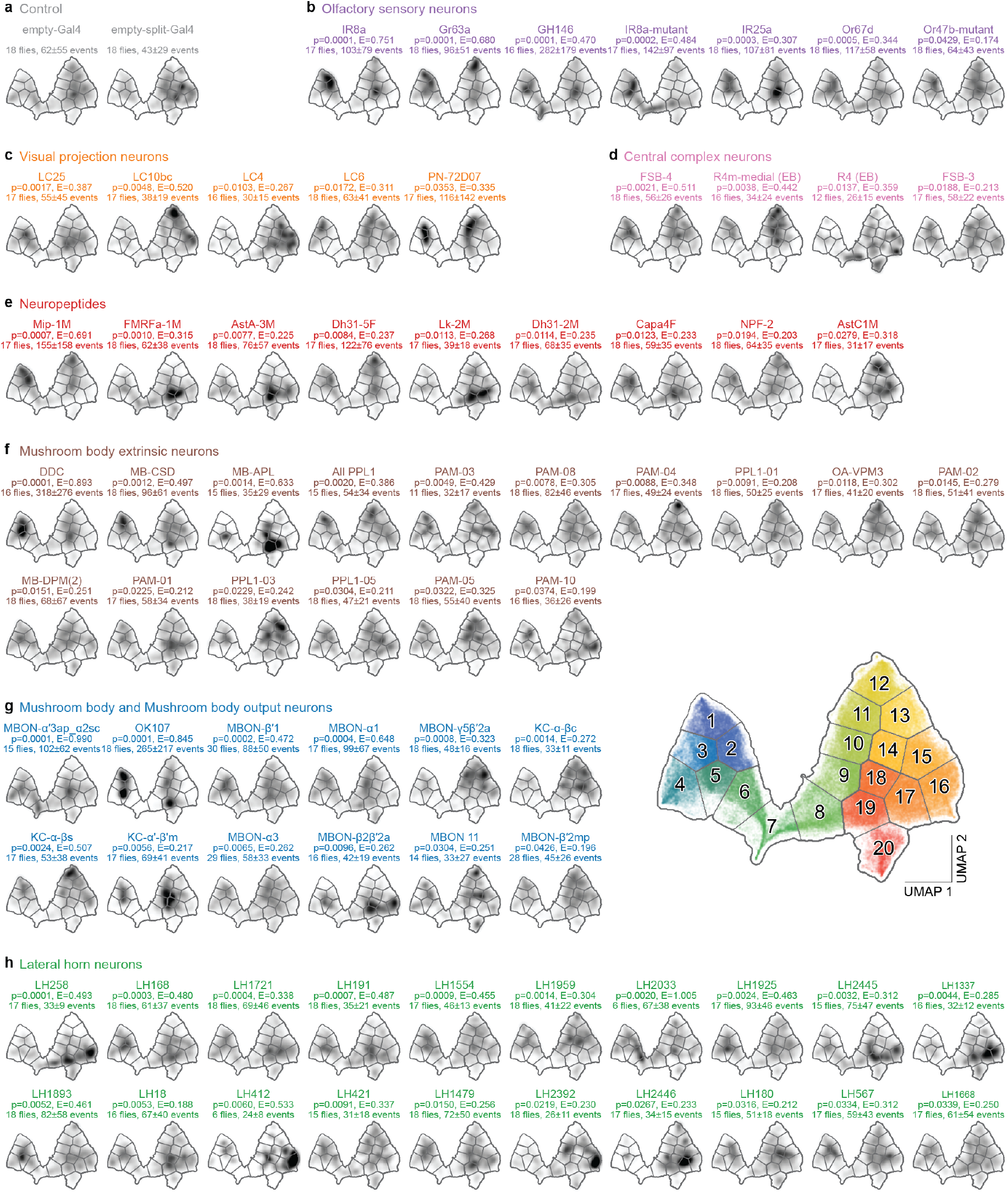
Driver lines with significantly different behavioral enrichment in UMAP space following neural silencing. UMAP density distributions of ball-contact events for driver lines, estimated by Gaussian kernel density estimation. Shading indicates local event density, with darker regions reflecting higher occupancy. P-values were computed by comparing each experimental line to its corresponding control using a permutation-based energy distance test (‘N’ indicates the number of animals represented). Gal4 driver lines were compared to the empty-Gal4 control; split-Gal4 driver lines were compared to the empty-split-Gal4 control, both shown in **(a)**. Only lines yielding a statistically significant difference in behavioral distributions are shown including lines for **(a)** controls, **(b)** olfactory sensory neurons, **(c)** visual projection neurons, **(d)** central complex neurons, **(e)** neuropeptidergic neurons, **(f)** mushroom body extrinsic neurons, **(g)** mushroom body intrinsic and output neurons, **(h)** lateral horn neurons.

**Extended Data Fig. 6:**
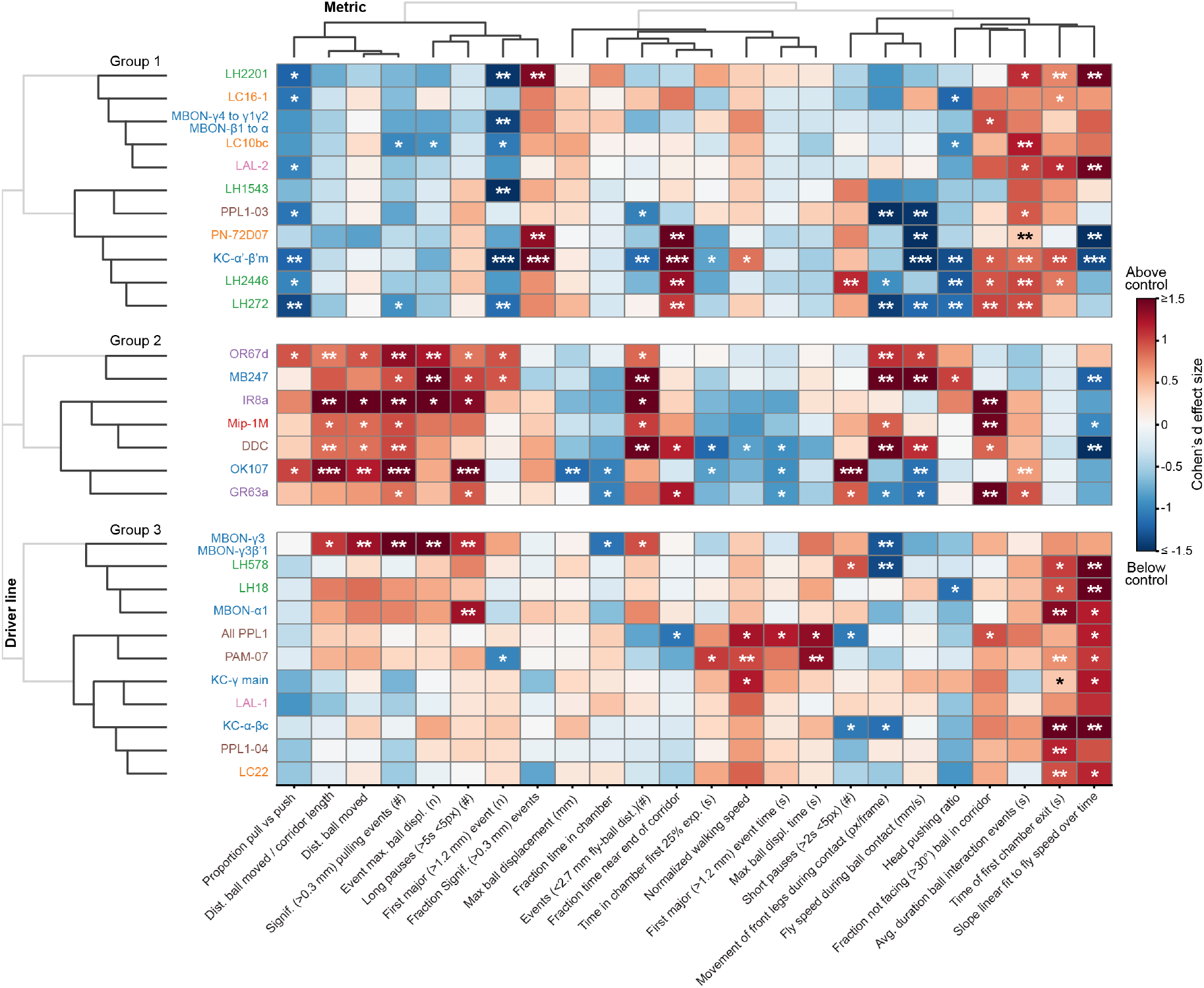
Hierarchical clustering of neural silencing screen high-consistency driver line hits and behavioral metrics. Heatmap showing ball pushing metrics for consistent screen hits (n = 29 driver lines with robust phenotypes, 18 flies per line). Metrics (columns) are hierarchically clustered by Pearson correlation distance. Driver lines (rows) are first grouped into 3 clusters by hierarchical clustering of Cohen’s d effect sizes (Euclidean distance, Ward linkage), then sorted within clusters. Colors represent Cohen’s d effect sizes compared with matched controls (empty-Gal4*>*UAS-TNT, empty-split-Gal4*>*UAS-TNT, or PR wild-type*>*UAS-TNT): dark blue (*d* ≤ −1.5) through white (no change) to dark red (*d* ≥ 1.5). Significance from permutation tests (FDR-corrected, *α* = 0.05) is indicated as: ns, p ≥ 0.05; *, p *<* 0.05; **, p *<* 0.01; ***, p *<* 0.001. Full statistical results are provided in Supplementary Data.

**Extended Data Fig. 7:**
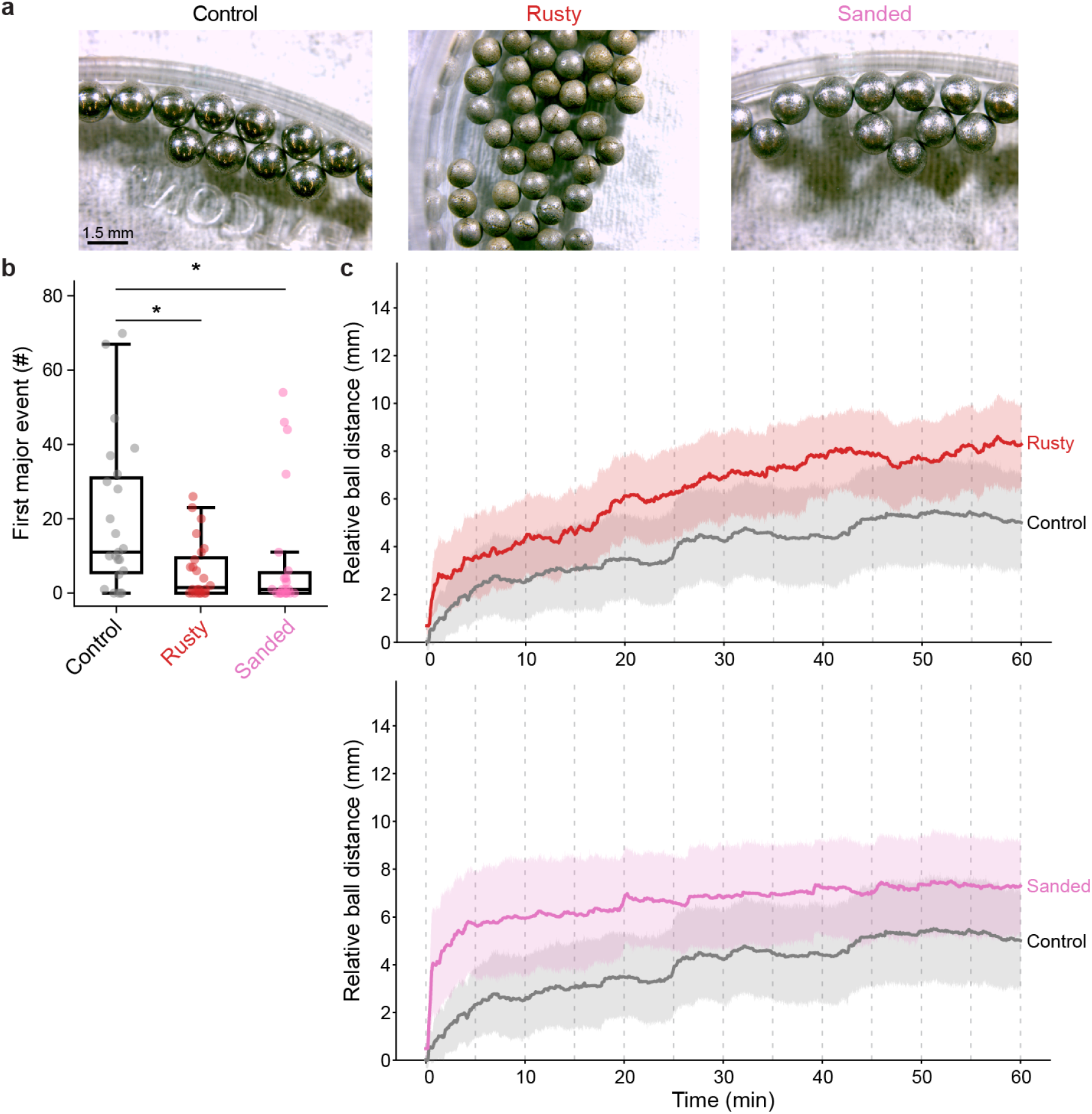
Impact of ball surface treatment on pushing. **(a)** Photos of balls for different surface treatments including untreated stainless steel balls (left), rusted by immersion in 90% ethanol for 24 h (middle), or sanded using P40 sandpaper (right). **(b)** Time to first major push (*>*1.2 mm) grouped as a function of treatment condition. Data shown as boxplots. Groups compared using permutation tests (n = 24 per group). Animals with either rusty or sanded balls both had a significantly earlier first major push compared to animals with untreated balls (rusted: *p* = 0.036; sanded: *p* = 0.036; permutation tests, FDR-corrected). Boxplots show the median (center line), interquartile range (IQR; box edges), 1.5 × IQR (whiskers), and individual data points (dots). **(c)** Distance the ball was moved relative to the start position for (top) control vs. rusty or (bottom) control vs. sanded balls. Shown are group averages with bootstrapped 95% CI in which data were binned into 5 min segments and compared using permutation tests with FDR correction (*α* = 0.05). No time bins reached significance after FDR correction for either treatment (all FDR-corrected *p* ≥ 0.08). Significance levels: ns, *p* ≥ 0.05; *, *p <* 0.05; **, *p <* 0.01; ***, *p <* 0.001.

**Extended Data Fig. 8:**
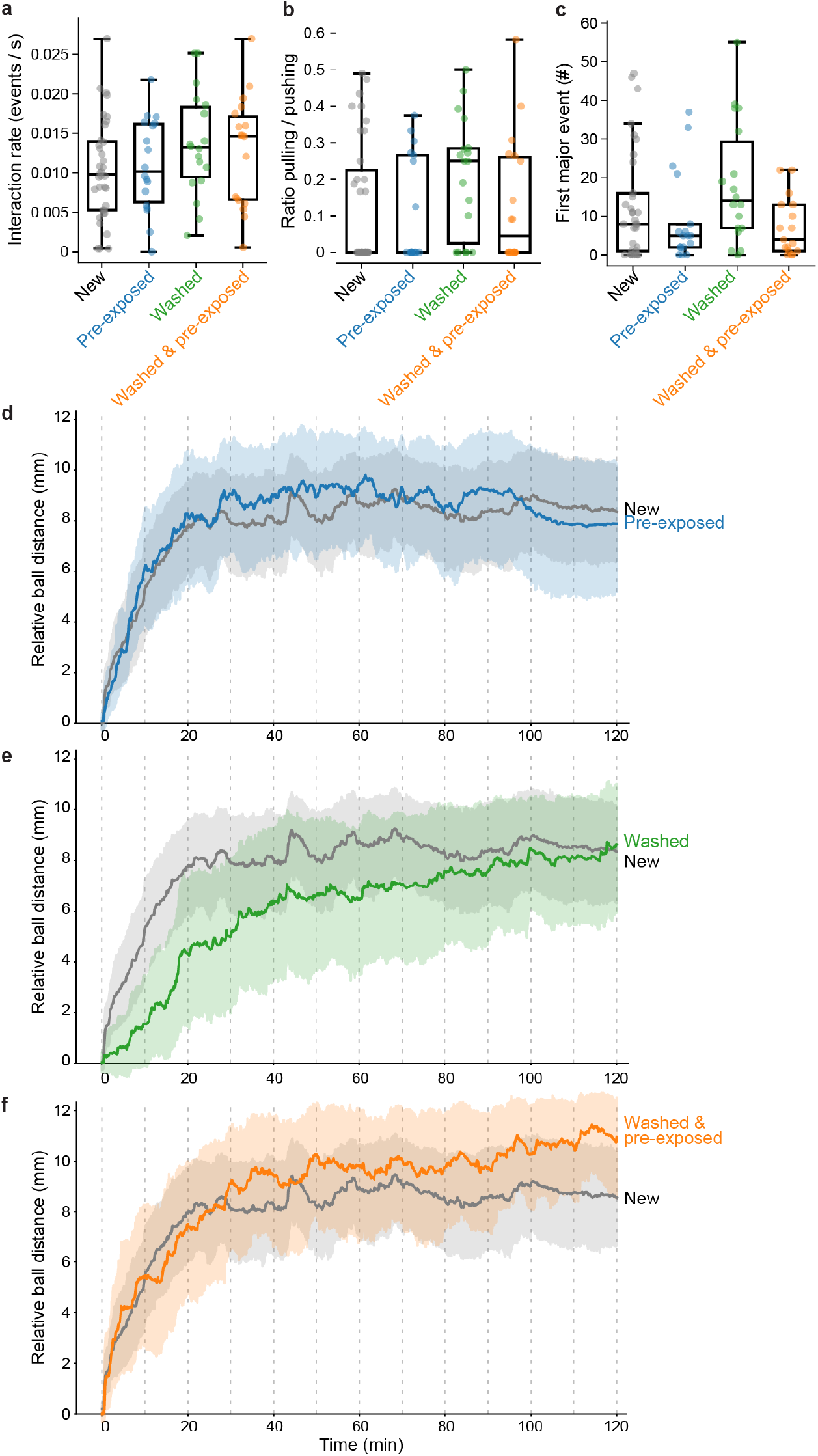
Effect of ball scent treatment on pushing. **(a-c)** Three behavioral metrics as a function of ball scent treatment: **(a)** interaction rate, **(b)** ratio of pulling /pushing significant interactions, and **(c)** the time until the first major push (*>*1.2 mm). Data shown as boxplots. Treatments are color-coded as new (gray), pre-exposed to fly odors (blue), washed with ethanol (green), or first washed and then pre-exposed (orange). All treatments were compared to control (‘new’) using pairwise permutation tests, FDR-corrected (*α* = 0.05) with n = 36 for control, and n = 17–18 for each treatment. No treatment produced a significant difference in any metric (all FDR-corrected *p* ≥ 0.13). Boxplots show the median (center line), interquartile range (IQR; box edges), 1.5 × IQR (whiskers), and individual data points (dots). **(d–f)** Distance the ball was moved relative to its start position for: **(d)** control vs pre-exposed to fly odors, **(e)** control vs ethanol washed, and **(f)** control vs washed and pre-exposed. (n = 36 for control, n = 18 per treatment). Data shown as group averages with bootstrapped 95% CI, binned into 10 min. segments and compared using permutation tests with FDR correction (*α* = 0.05). No time bin reached significance after FDR correction for any treatment (all FDR-corrected *p* ≥ 0.11). Significance levels: ns, *p* ≥ 0.05; *, *p <* 0.05; **, *p <* 0.01; ***, *p <* 0.001.

**Extended Data Fig. 9:**
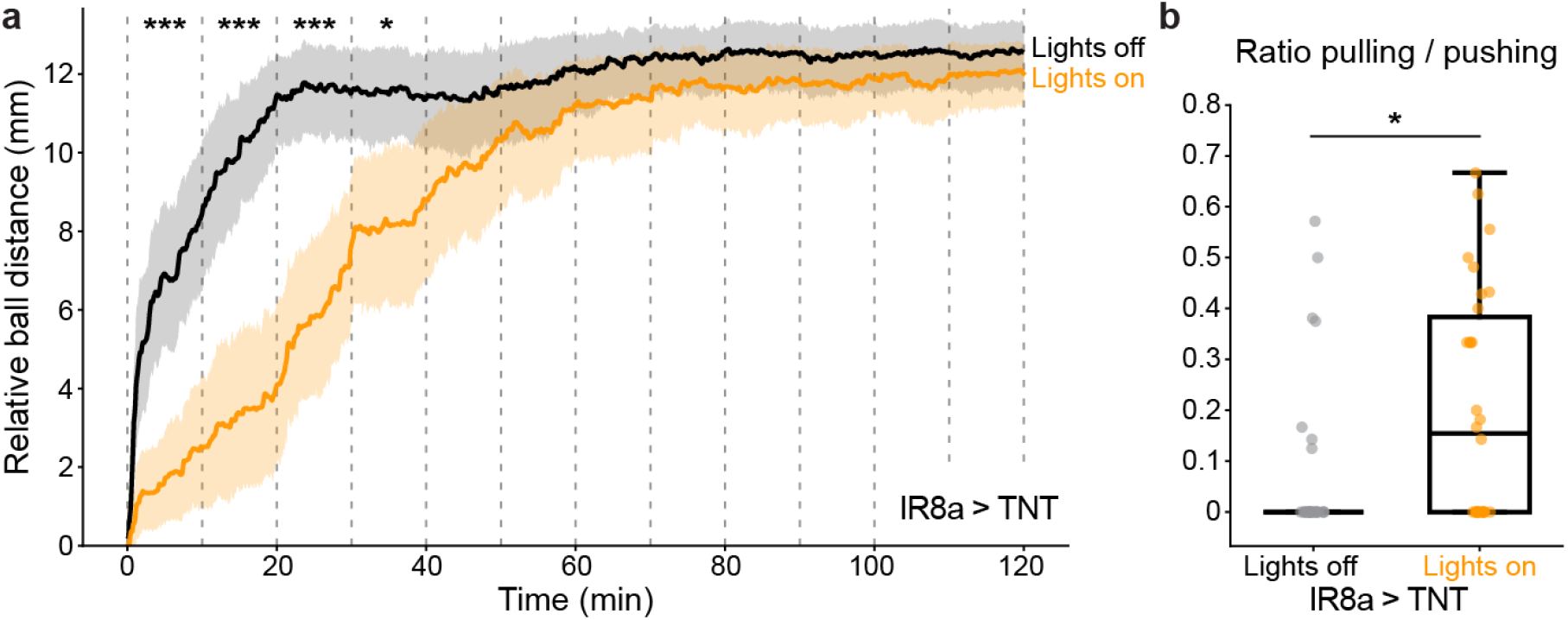
Ball pushing of IR8a-silenced animals under different illumination conditions. **(a)** Distance the ball was pushed over 2 h of the experiment relative to the ball’s start position. Shown are data either with (orange, n = 30 flies) or without (black, n = 30 flies) illumination plotted as means and bootstrapped 95% confidence intervals. Time-series data were binned into 10 min. segments and compared using permutation tests with FDR correction (*α* = 0.05). Animals in an illuminated arena pushed the ball significantly less in the first 40 minutes (bins 1–4: FDR-corrected *p <* 0.001; bin 3: *p* = 0.015), but not thereafter (bins 5–12: all *p* ≥ 0.23). **(b)** Ratio of pulling versus pushing significant events for lights off (gray, n = 30) or on (orange, n = 30 flies). Animals in an illuminated arena had a significantly higher ration of pulling versus pushing (permutation test, *p* = 0.018). Boxplots show the median (center line), interquartile range (IQR; box edges), 1.5 × IQR (whiskers), and individual data points (dots). Significance levels: ns, *p* ≥ 0.05; *, *p <* 0.05; **, *p <* 0.01; ***, *p <* 0.001.

**Extended Data Fig. 10:**
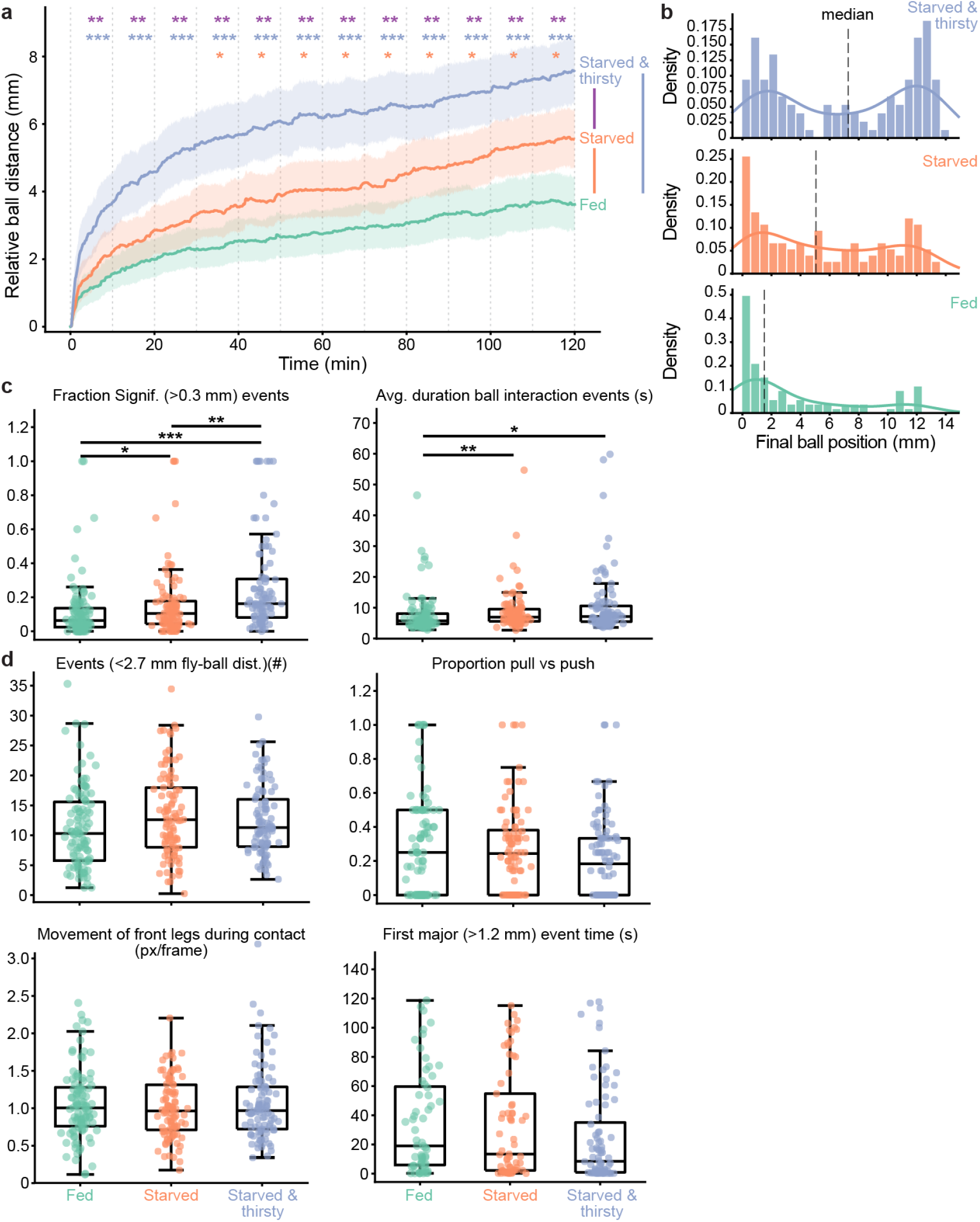
Effect of feeding state on ball manipulation. **(a)** Distance the ball was moved relative to its start position grouped by feeding state: fed (green, n = 84), starved (orange, n = 119), or starved and water deprived (blue, n = 119). Shown are group means (solid lines) with bootstrapped 95% confidence intervals (shading). Group data were binned into 5 min segments and compared using pairwise permutation tests with FDR correction (*α* = 0.05). Starved animals pushed significantly farther than fed animals from bin 4 onward (30–120 min; all FDRcorrected *p* ≤ 0.035), but not in bins 1–3 (all *p* ≥ 0.081). Starved and water-deprived animals pushed significantly farther than fed animals across all bins (all FDR-corrected *p <* 0.001), and significantly farther than starved animals across all bins (all FDR-corrected *p* ≤ 0.009). **(b)** Distribution of final ball positions relative to the start position as a function of feeding state (color-coded as in panel a). Histograms are overlaid with kernel density estimates. Median values are indicated by dashed vertical lines. **(c)** Behavioral metrics showing significant differences across feeding states: the ratio of significant interaction events (permutation tests, FDR-corrected: fed vs. starved *p* = 0.027; fed vs. starved and water-deprived *p <* 0.001; starved vs. starved and water-deprived *p* = 0.006) and interaction persistence (fed vs. starved *p* = 0.006; fed vs. starved and water-deprived *p* = 0.013; starved vs. starved and water-deprived ns). **(d)** Behavioral metrics showing no significant differences across nutritional states, including number of interaction events, pulling ratio, flailing, and time to first major push (all FDR-corrected *p* ≥ 0.16). Boxplots show the median (center line), interquartile range (IQR; box edges), 1.5 × IQR (whiskers), and individual data points (dots). Sample sizes per group for behavioral metrics (fed / starved / starved and water-deprived): significant ratio and interaction persistence: 102 / 101 / 101; number of events, pulling ratio, and flailing: 102 / 101 /101 (86–102 / 85–101 / 79–101 for metrics with stricter inclusion criteria); time to first major push: 60 / 69 / 84. Significance levels: ns, *p* ≥ 0.05; *, *p <* 0.05; **, *p <* 0.01; ***, *p <* 0.001.

## Supporting Information

### Supporting Information Table 1 - *Drosophila* transgenic driver lines

An Excel spreadsheet listing all transgenic driver lines used in this study. For each line, the Bloomington Drosophila Stock Center (BDSC) or Vienna Drosophila Resource Center (VDRC) stock number is indicated alongside the full genotype and a short name used throughout the manuscript. Control and genetic tool lines include a brief functional description, while screen driver lines are annotated with the targeted brain region or functional class.

Link to Supporting Information Table 1

### Supporting Information File 2 - Neural silencing screen: UMAP results for all driver lines

This file provides UMAP density maps of fly-ball contact event kinematics for all driver lines tested in the neural silencing screen, organized across three sub-files by line type. Each panel shows the density of contact events projected onto the 20-cluster UMAP space, with darker regions indicating higher event density. Each line is labeled with its identifier, the FDR-corrected permutation test *p*-value (*p*) and effect size (*E*) comparing its UMAP distribution to the matched control, and the number of flies and total contact events recorded. Lines are sorted by ascending *p*-value. Sub-file A contains results for 152 split-Gal4 lines, compared against the empty-split-Gal4 control (18 flies, 43 ± 25 events). Sub-file B contains results for 57 Gal4 lines, compared against the empty-Gal4 control (18 flies, 62 ± 55 events). Sub-file C contains results for 7 olfactory receptor mutant lines and two wild-type controls (TNT × PR and TNT × CS), displayed for reference.

Link to Supporting Information File 2

### Supporting Information File 3 - Neural silencing screen: ball pushing time course for all driver lines

Ball pushing time courses for all 212 driver lines, organized by targeted brain region. Each panel shows the mean ball position relative to the corridor start (solid lines) with bootstrapped 95% confidence intervals (shading) for a given experimental genotype (colored) and its matched control (gray; empty-Gal4*>*TNT, empty-split-Gal4*>*TNT, or PR*>*TNT). Data are binned into 10 min. segments over the 60 min. recording window. Experimental and control groups were compared at each time bin using permutation tests with FDR correction (*α* = 0.05); significant time bins are indicated by colored markers above each panel. Brain regions shown are: visual projection neurons, lateral horn neurons, central complex neurons, peripheral olfactory sensory neurons, mushroom body intrinsic neurons, mushroom body extrinsic and neuromodulatory neurons and neuropeptidergic neurons.

Link to Supporting Information File 3

### Supporting Information File 4 - Behavioral metrics for all 212 driver lines tested in the neural silencing screen

Heatmaps of Cohen’s *d* effect sizes for all driver lines tested in the silencing screen, computed relative to matched controls (empty-Gal4*>*TNT for simple Gal4 lines, empty-split-Gal4*>*TNT for split-Gal4 lines, or PR*>*TNT wild-type for mutant lines). Driver lines (rows) are ordered alphabetically within each brain region; Metrics (columns) are fixed in thematic groups. Cell colors represent Cohen’s *d* effect sizes clipped to [−1.5, 1.5], ranging from dark blue (*d* = − 1.5) through white (no change) to dark red (*d* = 1.5). Significance from permutation tests (10,000 iterations, FDR-corrected, *α* = 0.05) is indicated by asterisks (\**p <* 0.05, \*\**p <* 0.01, \*\*\**p <* 0.001).

Link to Supporting Information File 4

### Supporting Information File 5 - Neural silencing screen: consistency scores for all driver lines

Consistency scores across PCA configurations for all 212 driver lines tested in the neural silencing screen. Each bar indicates the proportion of the 20 PCA configurations (derived from a grid search over metric subsets, number of components, and, for Sparse PCA, sparsity regularization and ridge penalty) yielding a significant behavioral difference from matched controls (empty-Gal4*>*TNT, empty-split-Gal4*>*TNT, or wild-type PR*>*TNT). A genotype was classified as a hit (dashed line, 80% threshold, i.e., ≥ 16/20 configurations) if the multivariate permutation test, Mahalanobis distance, and at least one univariate Mann–Whitney *U* test on individual PC scores were all significant (FDR-corrected, *α* = 0.05). Driver lines are color-coded by brain region or functional class. The full distribution across all 212 lines is shown here to allow assessment of near-miss lines and overall screen sensitivity.

Link to Supporting Information File 5

### Supporting Information File 6 - Detailed statistics

Complete statistical results for all quantitative comparisons reported in the main figures and supplementary figures. Each sheet corresponds to one panel or set of panels, named by figure and panel identifier. For each comparison the following are reported: the metric tested, the genotypes or conditions compared, sample sizes (*n*) for each group, group means and medians, the effect estimate (raw difference between group means), Cohen’s *d* effect size, 95% bootstrapped confidence intervals of the difference expressed both as raw values and percentage change, the permutation test *p*-value, and the FDR-corrected *p*-value (Benjamini–Hochberg). For repeated-measures comparisons, the Friedman test statistic and pairwise Wilcoxon signed-rank test results are reported instead. For the neural silencing screen, results are provided for all 29 driver lines identified as robust hits across all 24 continuous behavioral metrics.

Link to Supporting Information File 6

## Supplementary Videos

### Video 1: Recording and tracking fly-ball interactions

Recordings of wild-type animals at the start of ball manipulation experiments, just before and after gate opening. Individual corridors from multiple arenas and recordings are cropped and concatenated. Indicated are tracked keypoints: right and left leg tips (red and blue, respectively, with dark-to-light shading from front to rear) as well as head, thorax, abdomen, and wings (white).

Link to Supplementary Video 1

### Video 2: High-resolution example of initial fly-ball probing interactions

High-resolution recording of a typical sequence of early interactions of the fly with the ball. Fly-ball contacts are automatically detected, during which the spatial overlap of the fly with the ball (red) and the ball center (cyan) are overlaid. The time during the experiment is indicated (top-left).

Link to Supplementary Video 2

### Video 3: High-resolution example of pushing during later fly-ball interactions

High-resolution recording of a later interaction in which the fly pushes the ball. Fly-ball contacts are automatically detected, during which the spatial overlap of the fly with the ball (red) and the ball center (cyan) are overlaid. The time during the experiment and a manual classification of the push-type is indicated (top-left). This is the same animal and experiment as in **Video 2**.

Link to Supplementary Video 3

### Video 4: Ball pushing types

Representative high-resolution examples of the main pushing types observed during fly-ball interactions including (i) one-leg interactions (00:00–00:07), (ii) two-leg interactions (00:10–00:17), (iii) rearing (00:19–00:34), and (iv) head (then single-leg) interactions (00:37–00:51). Videos are played at 0.2x speed.

Link to Supplementary Video 4

### Video 5: A fly interacting with an immobilized ball

Recording of a fly encountering a ball that has been immobilized using a hidden magnet to prevent ball displacement despite fly contact. The magnet is concealed under a back-lighting diffuser.

Link to Supplementary Video 5

### Video 6: A fly that is able to see but not touch an immobilized ball

Recording of a fly in a corridor in which a transparent plastic window prevents physical contact with a magnetically immobilized ball. This serves as a control condition for the magnetic immobilization experimental condition shown in **Video 5**.

Link to Supplementary Video 6

### Video 7: A fly finding and interacting with a test ball

Recording of a fly interacting with the test ball in the rectangular arena during the second hour of an experiment. Just prior to this, the training ball (top-right) was pushed or magnetically displaced, and the gate preventing premature entry into the second arena was removed.

Link to Supplementary Video 7

### Video 8: Behavioral videos and kinematics for UMAP cluster 4

Grid video of detected ball contacts from cluster 4 of the low-dimensional UMAP embedding, enriched in ball pulling interactions. Tracked keypoints are overlaid: right and left leg tips in red and blue, respectively (dark-to-light shading from front to rear). The ball is annotated in cyan. Trial-averaged, time-locked metric traces are also shown (left).

Link to Supplementary Video 8

### Video 9: Behavioral videos and kinematics for UMAP cluster 16

Grid video of detected ball contacts from cluster 16 of the low-dimensional UMAP embedding, enriched in ball pushing interactions. Tracked keypoints are overlaid: right and left leg tips in red and blue, respectively (dark-to-light shading from front to rear). The ball is annotated in cyan. Trial-averaged, time-locked metric traces are also shown (left).

Link to Supplementary Video 9

### Video 10: Significant ball manipulation events for a control (empty-split-Gal4*>*UAS-TNT) animal

Significant contact events (ball displacement ≥ 0.3 mm) were detected and concatenated for a empty-split-Gal4*>*UAS-TNT fly. Bottom axis tick marks show the chronological occurrence of each contact event. Fly-ball contacts are automatically detected, during which the spatial overlap of the fly with the ball (red) and the ball center (cyan) are overlaid. The time during the experiment is indicated (top-left).

Link to Supplementary Video 10

### Video 11: High-resolution example of ball pushing by LC10bc- and LC16-silenced animals

Early ball interactions in flies with silenced visual projection neurons, LC10bc and LC16, are shown sequentially. Fly-ball contacts are automatically detected, during which the spatial overlap of the fly with the ball (red) and the ball center (cyan) are overlaid. The time during the experiment is indicated (top-left).

Link to Supplementary Video 11

### Video 12: High-resolution example of pulling by control (empty-split-Gal4), Gr63a-, and Or67d-silenced animals

Significant ball pulling events (ball displacement≤ − 0.3 mm) were detected and concatenated for control flies (empty-split*>*TNT) and those with olfactory sensory neurons silenced (Gr63a*>*TNT and Or67d*>*TNT). These are shown sequentially. Bottom axis tick marks show the chronological occurrence of each contact event. Fly-ball contacts are automatically detected, during which the spatial overlap of the fly with the ball (red) and the ball center (cyan) are overlaid. Link to Supplementary Video 12

## Acknowledgments

We thank S. Boy-Röttger for assistance in animal preparation; R. Benton for sharing *D. mel*. transgenic driver lines; the members of the Neuroengineering Laboratory for helpful discussions and comments on the manuscript. Stocks obtained from the Bloomington Drosophila Stock Center (NIH P40OD018537) were used in this study. MD acknowledges support from a Fyssen foundation post-doctoral fellowship. TKCL acknowledges support from the Croucher Foundation. VLR acknowledges support from the Mexican National Council for Science and Technology, CONACYT, under the grant number 709993. PR acknowledges support from an SNSF Project Grant (175667).

## Author Contributions

M.D. - Conceptualization, Methodology, Software, Validation, Formal Analysis, Investigation, Data Curation, Validation, Writing – Original Draft Preparation, Writing – Review & Editing, Visualization.

D.D. - Methodology, Software, Validation, Formal Analysis, Investigation, Validation, Writing – Original Draft Preparation, Writing – Review & Editing, Visualization.

T.K.L. - Methodology, Software, Validation, Formal Analysis, Investigation, Validation, Writing – Original Draft Preparation, Writing – Review & Editing, Visualization.

V.L.R. - Conceptualization, Methodology, Software, Writing – Review & Editing.

P.R. - Conceptualization, Methodology, Resources, Writing – Original Draft Preparation, Writing - Review & Editing, Supervision, Project Administration, Funding Acquisition.

## Ethical compliance

All experiments were performed in compliance with relevant national (Switzerland) and institutional (EPFL) ethical regulations.

## References

[1] Shoko Sugasawa, Barbara Webb, and Susan D Healy. Object manipulation without hands. Proceedings of the Royal Society B, 288(1947):20203184, 2021.

[2] Antonino Errante, Settimio Ziccarelli, Gloria Mingolla, and Leonardo Fogassi. Grasping and manipulation: neural bases and anatomical circuitry in humans. Neuroscience, 458:203–212, 2021.

[3] Peter Janssen and Hansjörg Scherberger. Visual guidance in control of grasping. Annual review of neuroscience, 38(1):69–86, 2015.

[4] Maria Soledad Esposito, Paolo Capelli, and Silvia Arber. Brainstem nucleus mdv mediates skilled forelimb motor tasks. Nature, 508(7496):351–356, 2014.

[5] Eiman Azim, Juan Jiang, Bror Alstermark, and Thomas M Jessell. Skilled reaching relies on a v2a propriospinal internal copy circuit. Nature, 508(7496):357–363, 2014.

[6] Christian Rutz, Barbara C Klump, Lisa Komarczyk, Rosanna Leighton, Joshua Kramer, Saskia Wischnewski, Shoko Sugasawa, Michael B Morrissey, Richard James, James JH St Clair, et al. Discovery of species-wide tool use in the hawaiian crow. Nature, 537(7620):403–407, 2016.

[7] Chao Wen, Cai Wang, Xiaoli Guo, Hongyu Li, Haijun Xiao, Junbao Wen, and Shikui Dong. Object use in insects. Insect Science, 31(4):1001–1014, 2024.

[8] Olli J Loukola, Cwyn Solvi, Louie Coscos, and Lars Chittka. Bumblebees show cognitive flexibility by improving on an observed complex behavior. Science, 355(6327):833–836, 2017.

[9] Emily Baird, Marcus J Byrne, Jochen Smolka, Eric J Warrant, and Marie Dacke. The dung beetle dance: an orientation behaviour? PloS one, 7(1):e30211, 2012.

[10] JJ Gibson. The theory of affordances. Perceiving, acting and knowing: Towards an ecological psychology/Erlbaum, 1977.

[11] Natsuki Yamanobe, Weiwei Wan, Ixchel G Ramirez-Alpizar, Damien Petit, Tokuo Tsuji, Shuichi Akizuki, Manabu Hashimoto, Kazuyuki Nagata, and Kensuke Harada. A brief review of affordance in robotic manipulation research. Advanced Robotics, 31(19–20):1086–1101, 2017.

[12] Lorenzo Jamone, Emre Ugur, Angelo Cangelosi, Luciano Fadiga, Alexandre Bernardino, Justus Piater, and José Santos-Victor. Affordances in psychology, neuroscience, and robotics: A survey. IEEE Transactions on Cognitive and Developmental Systems, 10(1):4–25, 2016.

[13] Sridhar Ravi, Tim Siesenop, Olivier Bertrand, Liang Li, Charlotte Doussot, William H Warren, Stacey A Combes, and Martin Egelhaaf. Bumblebees perceive the spatial layout of their environment in relation to their body size and form to minimize inflight collisions. Proceedings of the National Academy of Sciences, 117(49):31494–31499, 2020.

[14] Laure-Anne Poissonnier, Yannick Hartmann, and Tomer J Czaczkes. Ants combine object affordance with latent learning to make efficient foraging decisions. Proceedings of the National Academy of Sciences, 120(35):e2302654120, 2023.

[15] Barret D. Pfeiffer, Teri-T B. Ngo, Karen L. Hibbard, Christine Murphy, Arnim Jenett, James W. Truman, and Gerald M. Rubin. Refinement of Tools for Targeted Gene Expression in Drosophila. Genetics, 186(2):735–755, October 2010.

[16] Alexander Shakeel Bates, Jasper S Phelps, Minsu Kim, Helen H Yang, Arie Matsliah, Zaki Ajabi, Eric Perlman, Kevin M Delgado, Mohammed Abdal Monium Osman, Christopher K Salmon, et al. Distributed control circuits across a brain-and-cord connectome. bioRxiv, 2025.

[17] Stuart Berg, Isabella R Beckett, Marta Costa, Philipp Schlegel, Michal Januszewski, Elizabeth C Marin, Aljoscha Nern, Stephan Preibisch, Wei Qiu, Shin-ya Takemura, et al. Sexual dimorphism in the complete connectome of the drosophila male central nervous system. bioRxiv, pages 2025–10, 2025.

[18] Gaby Maimon, Andrew D Straw, and Michael H Dickinson. Active flight increases the gain of visual motion processing in drosophila. Nature neuroscience, 13(3):393–399, 2010.

[19] Johannes D Seelig, M Eugenia Chiappe, Gus K Lott, Anirban Dutta, Jason E Osborne, Michael B Reiser, and Vivek Jayaraman. Two-photon calcium imaging from head-fixed drosophila during optomotor walking behavior. Nature methods, 7(7):535–540, 2010.

[20] Chin-Lin Chen, Laura Hermans, Meera C. Viswanathan, Denis Fortun, Florian Aymanns, Michael Unser, Anthony Cammarato, Michael H. Dickinson, and Pavan Ramdya. Imaging neural activity in the ventral nerve cord of behaving adult Drosophila. Nature Communications, 9(1):4390, December 2018.

[21] Kenichi Iwasaki, Sena Kawano, Ashnu Cassod, Charles Neuhauser, and Aleksandr Rayshubskiy. Drosophila learn to prefer immobile spherical objects through repeated physical interaction. Current Biology, 2025.

[22] Alice A Robie, Andrew D Straw, and Michael H Dickinson. Object preference by walking fruit flies, drosophila melanogaster, is mediated by vision and graviperception. Journal of Experimental Biology, 213(14):2494–2506, 2010.

[23] Talmo D. Pereira, Nathaniel Tabris, Arie Matsliah, David M. Turner, Junyu Li, Shruthi Ravindranath, Eleni S. Papadoyannis, Edna Normand, David S. Deutsch, Z. Yan Wang, Grace C. McKenzie-Smith, Catalin C. Mitelut, Marielisa Diez Castro, John D’Uva, Mikhail Kislin, Dan H. Sanes, Sarah D. Kocher, Samuel S.-H. Wang, Annegret L. Falkner, Joshua W. Shaevitz, and Mala Murthy. SLEAP: A deep learning system for multi-animal pose tracking. Nature Methods, 19(4):486–495, April 2022.

[24] Ming Wu, Aljoscha Nern, W Ryan Williamson, Mai M Morimoto, Michael B Reiser, Gwyneth M Card, and Gerald M Rubin. Visual projection neurons in the drosophila lobula link feature detection to distinct behavioral programs. Elife, 5:e21022, 2016.

[25] Benjamin R Cowley, Adam J Calhoun, Nivedita Rangarajan, Elise Ireland, Maxwell H Turner, Jonathan W Pillow, and Mala Murthy. Mapping model units to visual neurons reveals population code for social behaviour. Nature, 629(8014):1100–1108, 2024.

[26] Rajyashree Sen, Ming Wu, Kristin Branson, Alice Robie, Gerald M. Rubin, and Barry J. Dickson. Moonwalker Descending Neurons Mediate Visually Evoked Retreat in Drosophila. Current Biology, 27(5):766–771, March 2017.

[27] Nathan C Klapoetke, Aljoscha Nern, Edward M Rogers, Gerald M Rubin, Michael B Reiser, and Gwyneth M Card. A functionally ordered visual feature map in the drosophila brain. Neuron, 110(10):1700–1711, 2022.

[28] Inês MA Ribeiro, Michael Drews, Armin Bahl, Christian Machacek, Alexander Borst, and Barry J Dickson. Visual projection neurons mediating directed courtship in drosophila. Cell, 174(3):607–621, 2018.

[29] Sudeshna Das Chakraborty and Silke Sachse. Olfactory processing in the lateral horn of drosophila. Cell and Tissue Research, 383(1):113–123, 2021.

[30] Michael-John Dolan, Shahar Frechter, Alexander Shakeel Bates, Chuntao Dan, Paavo Huoviala, Ruairí Jv Roberts, Philipp Schlegel, Serene Dhawan, Remy Tabano, Heather Dionne, et al. Neurogenetic dissection of the drosophila lateral horn reveals major outputs, diverse behavioural functions, and interactions with the mushroom body. Elife, 8:e43079, 2019.

[31] Shahar Frechter, Alexander Shakeel Bates, Sina Tootoonian, Michael-John Dolan, James Manton, Arian Rokkum Jamasb, Johannes Kohl, Davi Bock, and Gregory Jefferis. Functional and anatomical specificity in a higher olfactory centre. elife, 8:e44590, 2019.

[32] Soohong Min, Hyo-Seok Chae, Yong-Hoon Jang, Sekyu Choi, Sion Lee, Yong Taek Jeong, Walton D Jones, Seok Jun Moon, Young-Joon Kim, and Jongkyeong Chung. Identification of a peptidergic pathway critical to satiety responses in drosophila. Current Biology, 26(6):814–820, 2016.

[33] Ashiq Hussain, Habibe K Ü çpunar, Mo Zhang, Laura F Loschek, and Ilona C Grunwald Kadow. Neuropeptides modulate female chemosensory processing upon mating in drosophila. PLoS biology, 14(5):e1002455, 2016.

[34] Tyler R Sizemore, Julius Jonaitis, and Andrew M Dacks. Heterogeneous receptor expression underlies non-uniform peptidergic modulation of olfaction in drosophila. Nature Communications, 14(1):5280, 2023.

[35] Jae Young Kwon, Anupama Dahanukar, Linnea A Weiss, and John R Carlson. The molecular basis of co2 reception in drosophila. Proceedings of the National Academy of Sciences, 104(9):3574–3578, 2007.

[36] Greg SB Suh, Allan M Wong, Anne C Hergarden, Jing W Wang, Anne F Simon, Seymour Benzer, Richard Axel, and David J Anderson. A single population of olfactory sensory neurons mediates an innate avoidance behaviour in drosophila. Nature, 431(7010):854–859, 2004.

[37] Amina Kurtovic, Alexandre Widmer, and Barry J Dickson. A single class of olfactory neurons mediates behavioural responses to a drosophila sex pheromone. Nature, 446(7135):542–546, 2007.

[38] Liming Wang and David J Anderson. Identification of an aggression-promoting pheromone and its receptor neurons in drosophila. Nature, 463(7278):227–231, 2010.

[39] Meghan Laturney and Jean-Christophe Billeter. Drosophila melanogaster females restore their attractiveness after mating by removing male anti-aphrodisiac pheromones. Nature communications, 7(1):12322, 2016.

[40] Scott E Dobrin and Susan E Fahrbach. Visual associative learning in restrained honey bees with intact antennae. PloS one, 7(6):e37666, 2012.

[41] Takeshi Morita, Nia G Lyn, Ricarda K von Heynitz, Olivia V Goldman, Trevor R Sorrells, Matthew DeGennaro, Benjamin J Matthews, Leah Houri-Zeevi, and Leslie B Vosshall. Cross-modal sensory compensation increases mosquito attraction to humans. Science Advances, 11(1):eadn5758, 2025.

[42] Pierre Junca, Molly Stanley, Pierre-Yves Musso, and Michael D Gordon. Modulation of taste sensitivity by the olfactory system in drosophila. bioRxiv, pages 2021–03, 2021.

[43] Pierre-Yves Oudeyer and Frederic Kaplan. What is intrinsic motivation? a typology of computational approaches. Frontiers in neurorobotics, 1:108, 2007.

[44] William G Quinn and Ralph J Greenspan. Learning and courtship in drosophila: two stories with mutants. Annual review of neuroscience, 1984.

[45] Margaret S Livingstone and Bruce L Tempel. Genetic dissection of monoamine neurotransmitter synthesis in drosophila. Nature, 303(5912):67–70, 1983.

[46] Mehrab N Modi, Yichun Shuai, and Glenn C Turner. The drosophila mushroom body: from architecture to algorithm in a learning circuit. Annual review of neuroscience, 43(1):465–484, 2020.

[47] Johannes Felsenberg. Changing memories on the fly: the neural circuits of memory re-evaluation in drosophila melanogaster. Current opinion in neurobiology, 67:190–198, 2021.

[48] Yvette E Fisher, Michael Marquis, Isabel D’Alessandro, and Rachel I Wilson. Dopamine promotes head direction plasticity during orienting movements. Nature, 612(7939):316–322, 2022.

[49] Ronald L Davis. Learning and memory using drosophila melanogaster: a focus on advances made in the fifth decade of research. Genetics, 224(4):iyad085, 2023.

[50] Yoshinori Aso, Divya Sitaraman, Toshiharu Ichinose, Karla R Kaun, Katrin Vogt, Ghislain Belliart-Guérin, Pierre-Yves Plaçais, Alice A Robie, Nobuhiro Yamagata, Christopher Schnaitmann, et al. Mushroom body output neurons encode valence and guide memory-based action selection in drosophila. elife, 3:e04580, 2014.

[51] Toshiharu Ichinose, Mai Kanno, Hongyang Wu, Nobuhiro Yamagata, Huan Sun, Ayako Abe, and Hiromu Tanimoto. Mushroom body output differentiates memory processes and distinct memory-guided behaviors. Current Biology, 31(6):1294–1302, 2021.

[52] Ulrike Pech, Shubham Dipt, Jonas Barth, Priyanka Singh, Mandy Jauch, Andreas S Thum, André Fiala and Thomas Riemensperger. Mushroom body miscellanea: transgenic drosophila strains expressing anatomical and physiological sensor proteins in kenyon cells. Frontiers in Neural Circuits, 7:147, 2013.

[53] Paul Cisek and Andrea M Green. Toward a neuroscience of natural behavior. Current opinion in Neurobiology, 86:102859, 2024.

[54] Pu Fan, Devanand S Manoli, Osama M Ahmed, Yi Chen, Neha Agarwal, Sara Kwong, Allen G Cai, Jeffrey Neitz, Adam Renslo, Bruce S Baker, et al. Genetic and neural mechanisms that inhibit drosophila from mating with other species. Cell, 154(1):89–102, 2013.

[55] John C. Tuthill and Rachel I. Wilson. Mechanosensation and Adaptive Motor Control in Insects. Current Biology, 26(20):R1022–R1038, October 2016.

[56] Shuke Xiao, Lisa S Baik, Xueying Shang, and John R Carlson. Meeting a threat of the anthropocene: taste avoidance of metal ions by drosophila. Proceedings of the National Academy of Sciences, 119(25):e2204238119, 2022.

[57] Chenghao Chen, Edgar Buhl, Min Xu, Vincent Croset, Johanna S Rees, Kathryn S Lilley, Richard Benton, James JL Hodge, and Ralf Stanewsky. Drosophila ionotropic receptor 25a mediates circadian clock resetting by temperature. Nature, 527(7579):516–520, 2015.

[58] Peter Redgrave and Kevin Gurney. The short-latency dopamine signal: a role in discovering novel actions? Nature reviews neuroscience, 7(12):967–975, 2006.

[59] Jeffrey E Markowitz, Winthrop F Gillis, Maya Jay, Jeffrey Wood, Ryley W Harris, Robert Cieszkowski, Rebecca Scott, David Brann, Dorothy Koveal, Tomasz Kula, et al. Spontaneous behaviour is structured by reinforcement without explicit reward. Nature, 614(7946):108–117, 2023.

[60] Vijay MK Namboodiri. “but why?” dopamine and causal learning. Current Opinion in Behavioral Sciences, 60:101443, 2024.

[61] Huijeong Jeong, Annie Taylor, Joseph R Floeder, Martin Lohmann, Stefan Mihalas, Brenda Wu, Mingkang Zhou, Dennis A Burke, and Vijay Mohan K Namboodiri. Mesolimbic dopamine release conveys causal associations. Science, 378(6626):eabq6740, 2022.

[62] Sophie Aimon, Karen Y Cheng, Julijana Gjorgjieva, and Ilona C Grunwald Kadow. Global change in brain state during spontaneous and forced walk in drosophila is composed of combined activity patterns of different neuron classes. Elife, 12:e85202, 2023.

[63] Fatima Amin, Jasmine T Stone, Christian König, Nino Mancini, Kazuma Murakami, Salil S Bidaye, M-Marcel Heim, David Owald, Utsab Majumder, Ilona C Grunwald Kadow, et al. Avoidance engages dopaminergic punishment in drosophila. bioRxiv, 2025.

[64] Korleki Akiti, Iku Tsutsui-Kimura, Yudi Xie, Alexander Mathis, Jeffrey E Markowitz, Rockwell Anyoha, Sandeep Robert Datta, Mackenzie Weygandt Mathis, Naoshige Uchida, and Mitsuko Watabe-Uchida. Striatal dopamine explains novelty-induced behavioral dynamics and individual variability in threat prediction. Neuron, 110(22):3789–3804, 2022.

[65] Kay C Montgomery. The role of the exploratory drive in learning. Journal of Comparative and Physiological Psychology, 47(1):60, 1954.

[66] Harry F Harlow, Margaret Kuenne Harlow, and Donald R Meyer. Learning motivated by a manipulation drive. Journal of experimental psychology, 40(2):228, 1950.

[67] Celeste Kidd and Benjamin Y Hayden. The psychology and neuroscience of curiosity. Neuron, 88(3):449–460, 2015.

[68] Daniel E Berlyne. Curiosity and exploration: Animals spend much of their time seeking stimuli whose significance raises problems for psychology. Science, 153(3731):25–33, 1966.

[69] Tilman Triphan, Clara H Ferreira, and Wolf Huetteroth. Play-like behavior exhibited by the vinegar fly drosophila melanogaster. Current Biology, 35(5):1145–1155, 2025.

[70] Hiruni Samadi Galpayage Dona, Cwyn Solvi, Amelia Kowalewska, Kaarle Mäkelä, HaDi MaBouDi, and Lars Chittka. Do bumble bees play? Animal Behaviour, 194:239–251, 2022.

[71] Gordon M Burghardt. The genesis of animal play: Testing the limits. MIT press, 2005.

[72] Ravi Allada and Brian Y Chung. Circadian organization of behavior and physiology in drosophila. Annual review of physiology, 72(1):605–624, 2010.

[73] Pavan Ramdya, Robin Thandiackal, Raphael Cherney, Thibault Asselborn, Richard Benton, Auke Jan Ijspeert, and Dario Floreano. Climbing favours the tripod gait over alternative faster insect gaits. Nature Communications, 8(1):1–11, February 2017.

[74] David Tadres and Matthieu Louis. Pivr: An affordable and versatile closed-loop platform to study unrestrained sensorimotor behavior. PLoS biology, 18(7):e3000712, 2020.

[75] Mark Sandler, Andrew Howard, Menglong Zhu, Andrey Zhmoginov, and Liang-Chieh Chen. Mobilenetv2: Inverted residuals and linear bottlenecks. In Proceedings of the IEEE conference on computer vision and pattern recognition, pages 4510–4520, 2018.

[76] Talmo D. Pereira, Joshua W. Shaevitz, and Mala Murthy. Quantifying behavior to understand the brain. Nature Neuroscience, 23(12):1537–1549, December 2020.

[77] Talmo D Pereira, Diego E Aldarondo, Lindsay Willmore, Mikhail Kislin, Samuel S-H Wang, Mala Murthy, and Joshua W Shaevitz. Fast animal pose estimation using deep neural networks. Nature methods, 16(1):117–125, 2019.

[78] John A Bogovic, Hideo Otsuna, Larissa Heinrich, Masayoshi Ito, Jennifer Jeter, Geoffrey Meissner, Aljoscha Nern, Jennifer Colonell, Oz Malkesman, Kei Ito, et al. An unbiased template of the drosophila brain and ventral nerve cord. Plos one, 15(12):e0236495, 2020.

