## Supplementary figures and images for "Object manipulation and affordance learning in *Drosophila*"

### Full_euclidean_distance_coordinates_line_Control.png

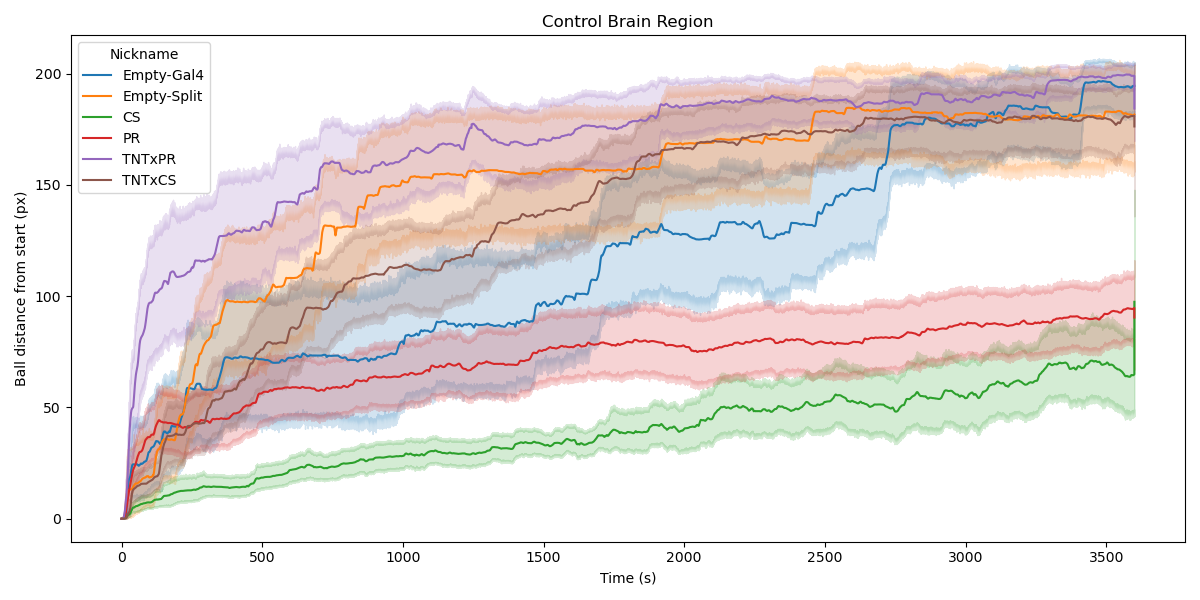

### Full_euclidean_distance_coordinates_line_CX.png

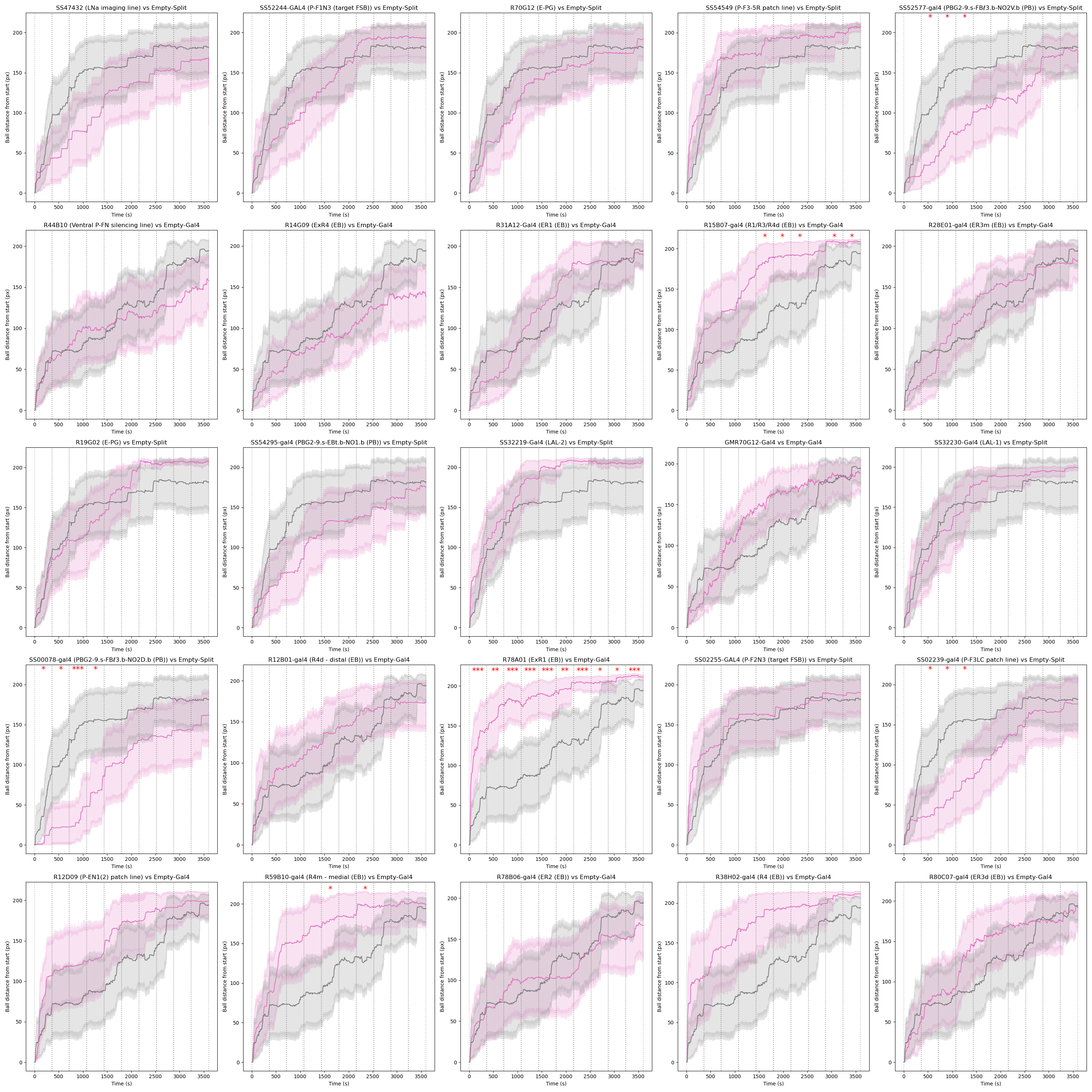

### Full_euclidean_distance_coordinates_line_LH.png

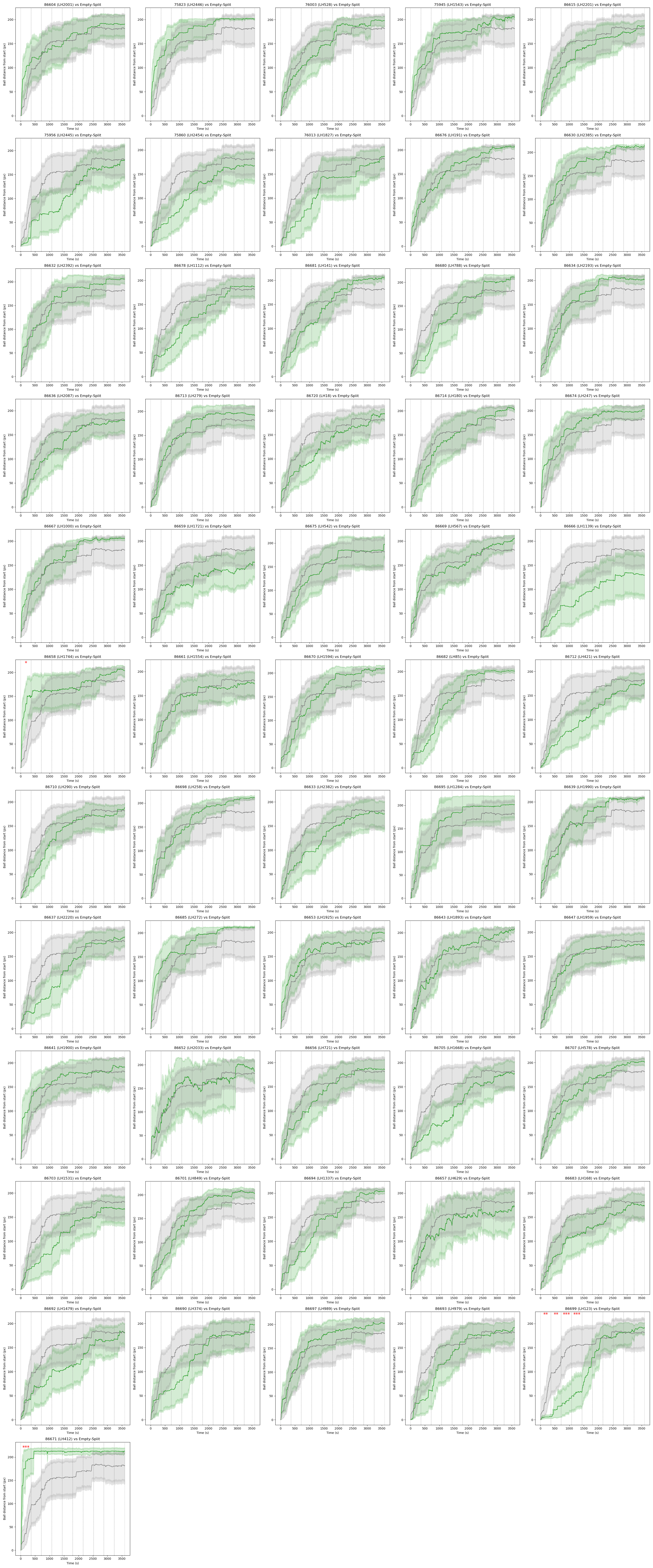

### Full_euclidean_distance_coordinates_line_MB extrinsic neurons.png

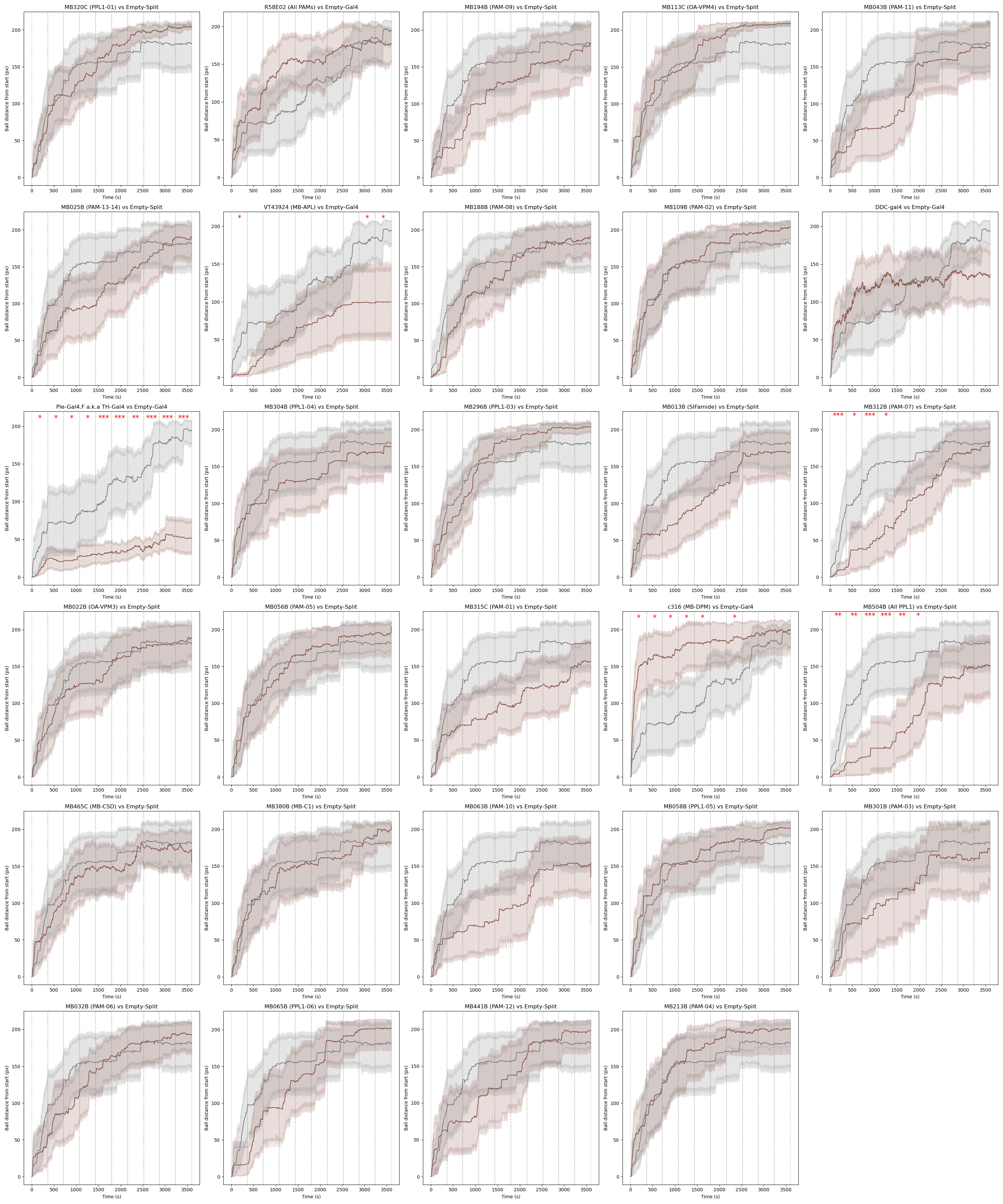

### Full_euclidean_distance_coordinates_line_MB.png

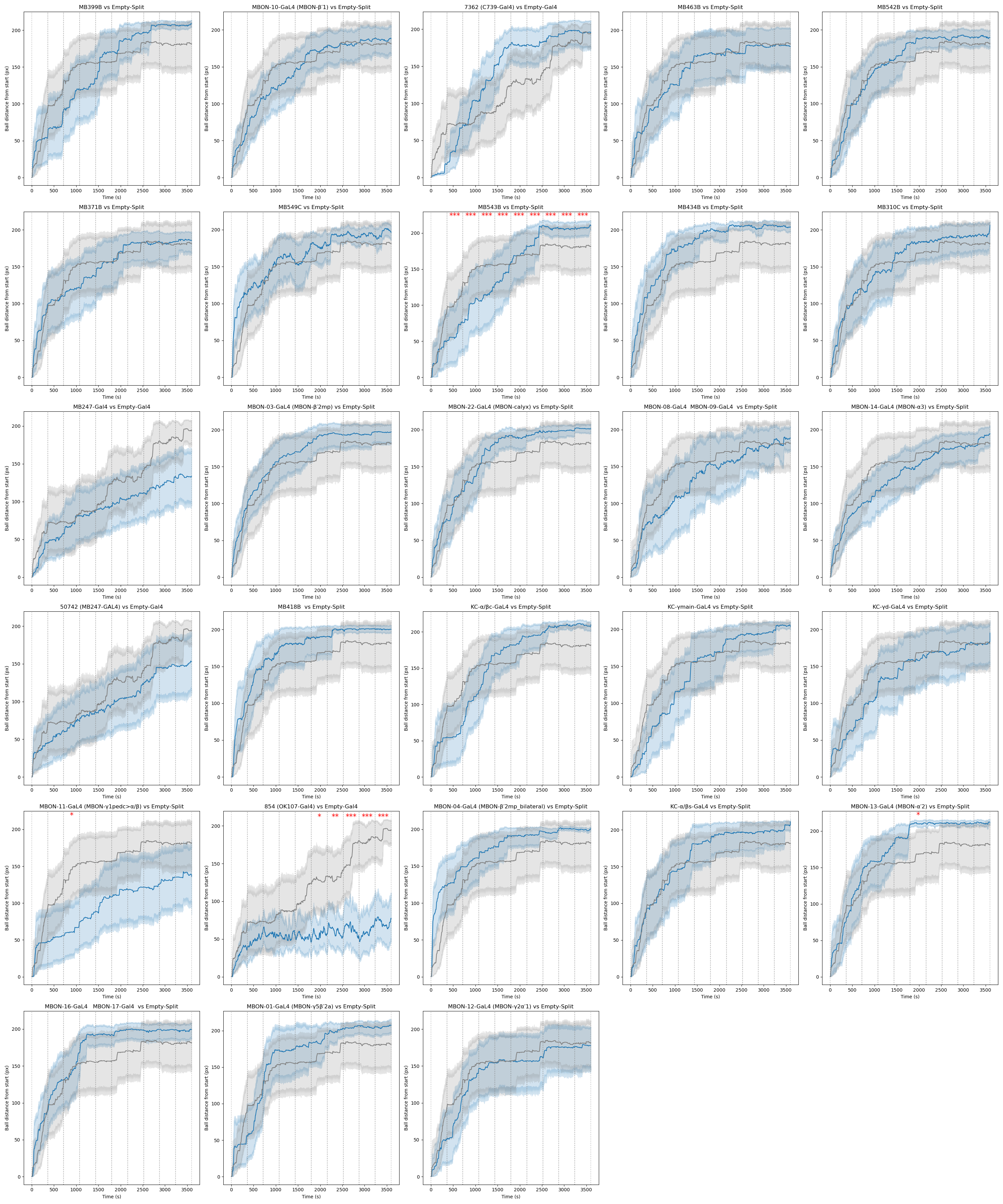

### Full_euclidean_distance_coordinates_line_Neuropeptide.png

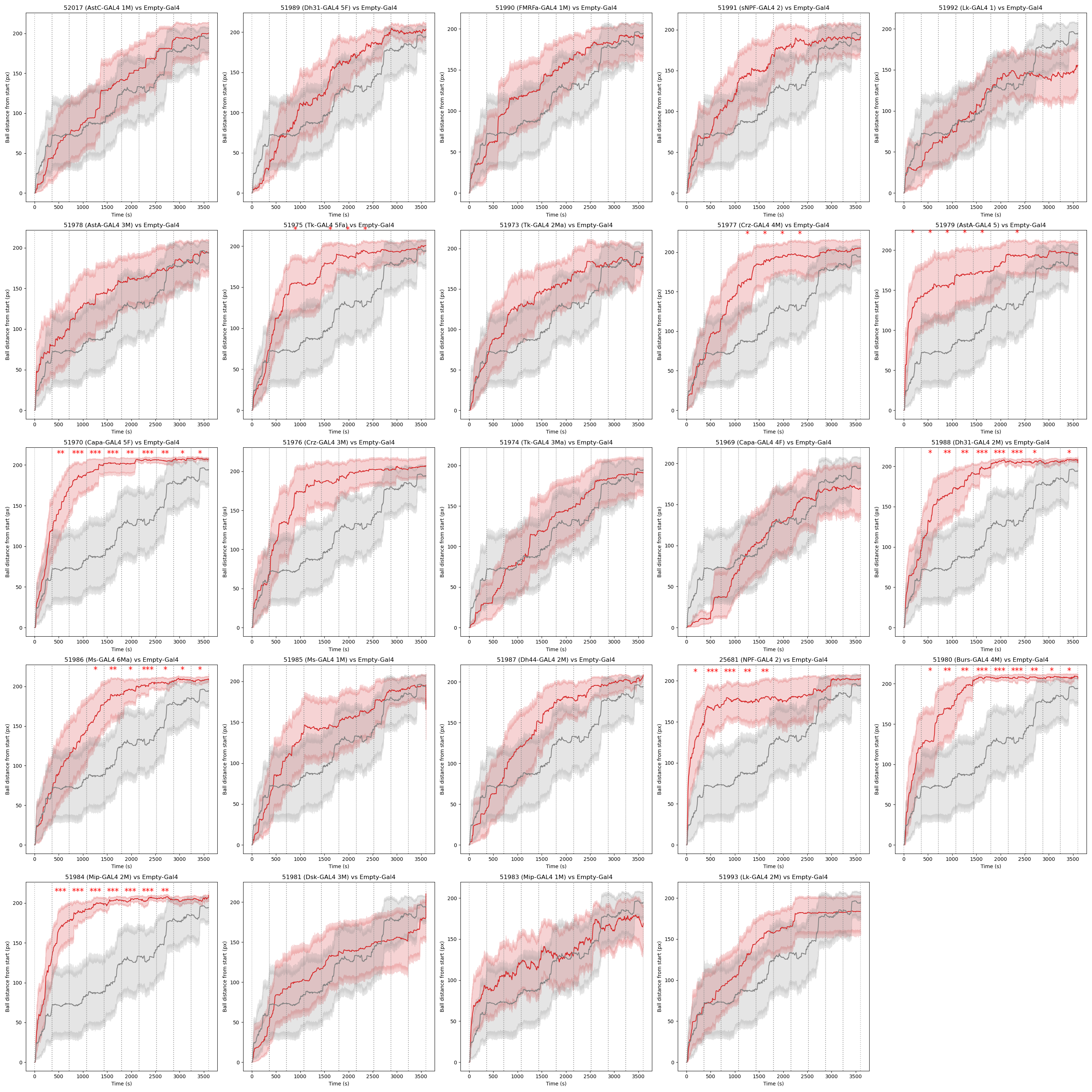

### Full_euclidean_distance_coordinates_line_Olfaction.png

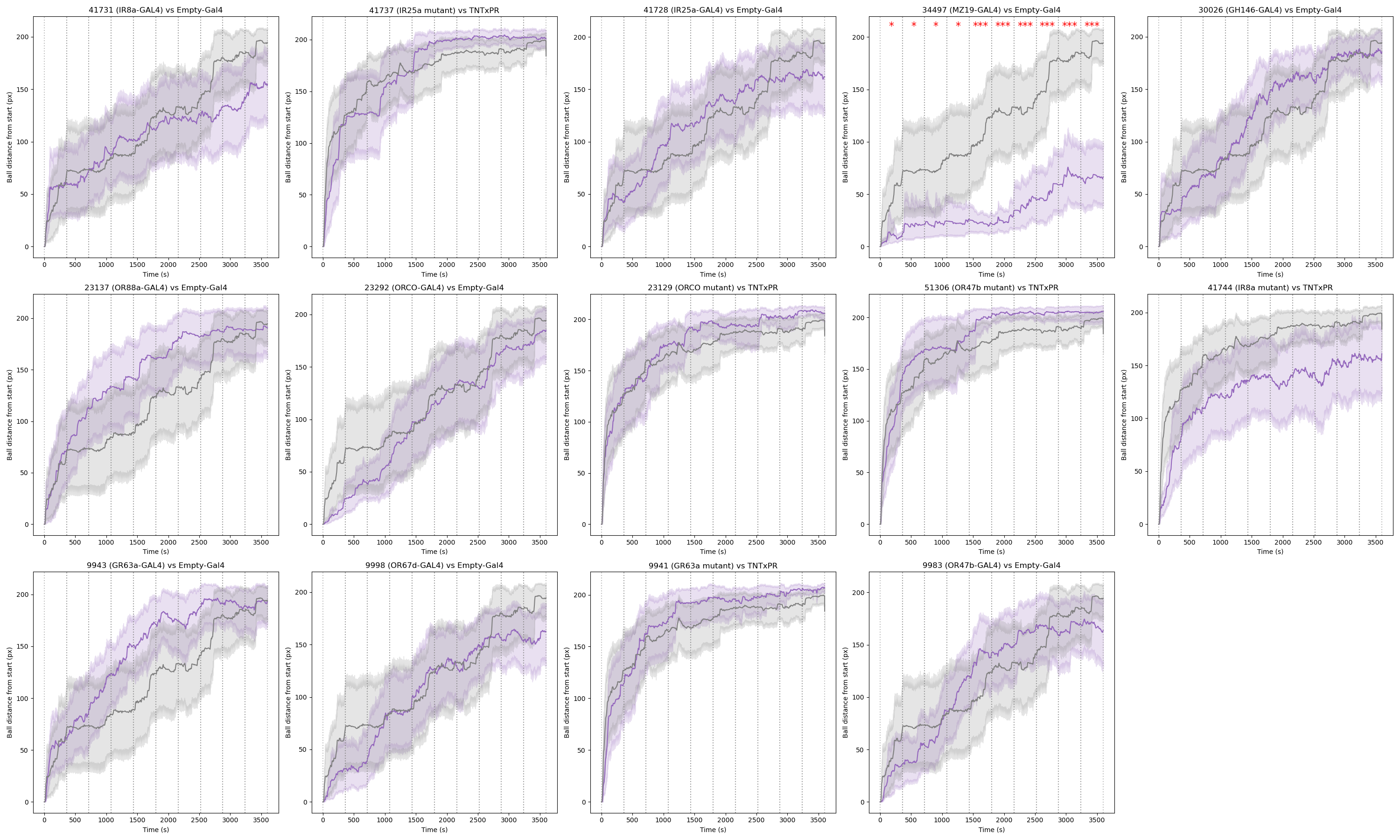

### Full_euclidean_distance_coordinates_line_Vision.png

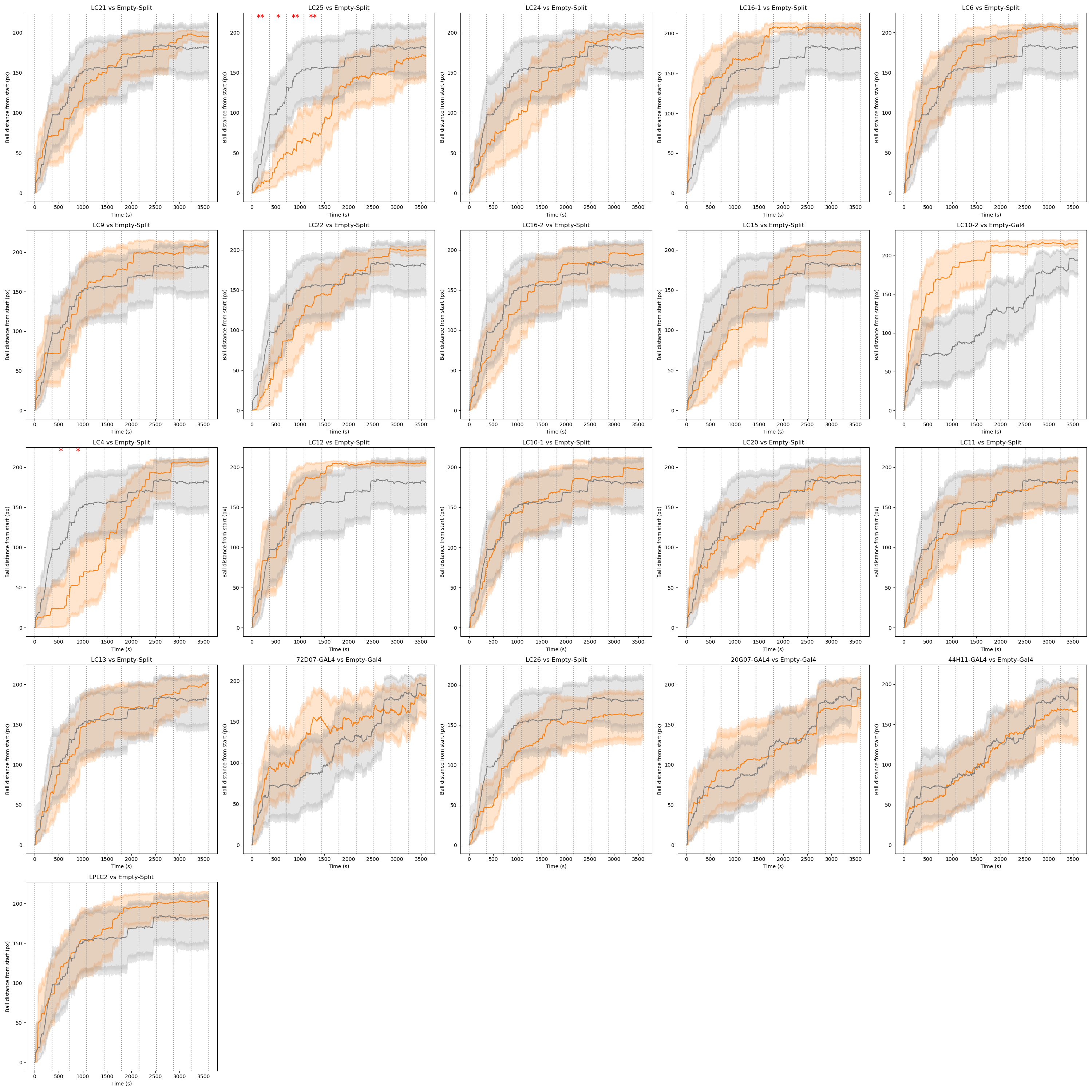

### lines_gal4.pdf

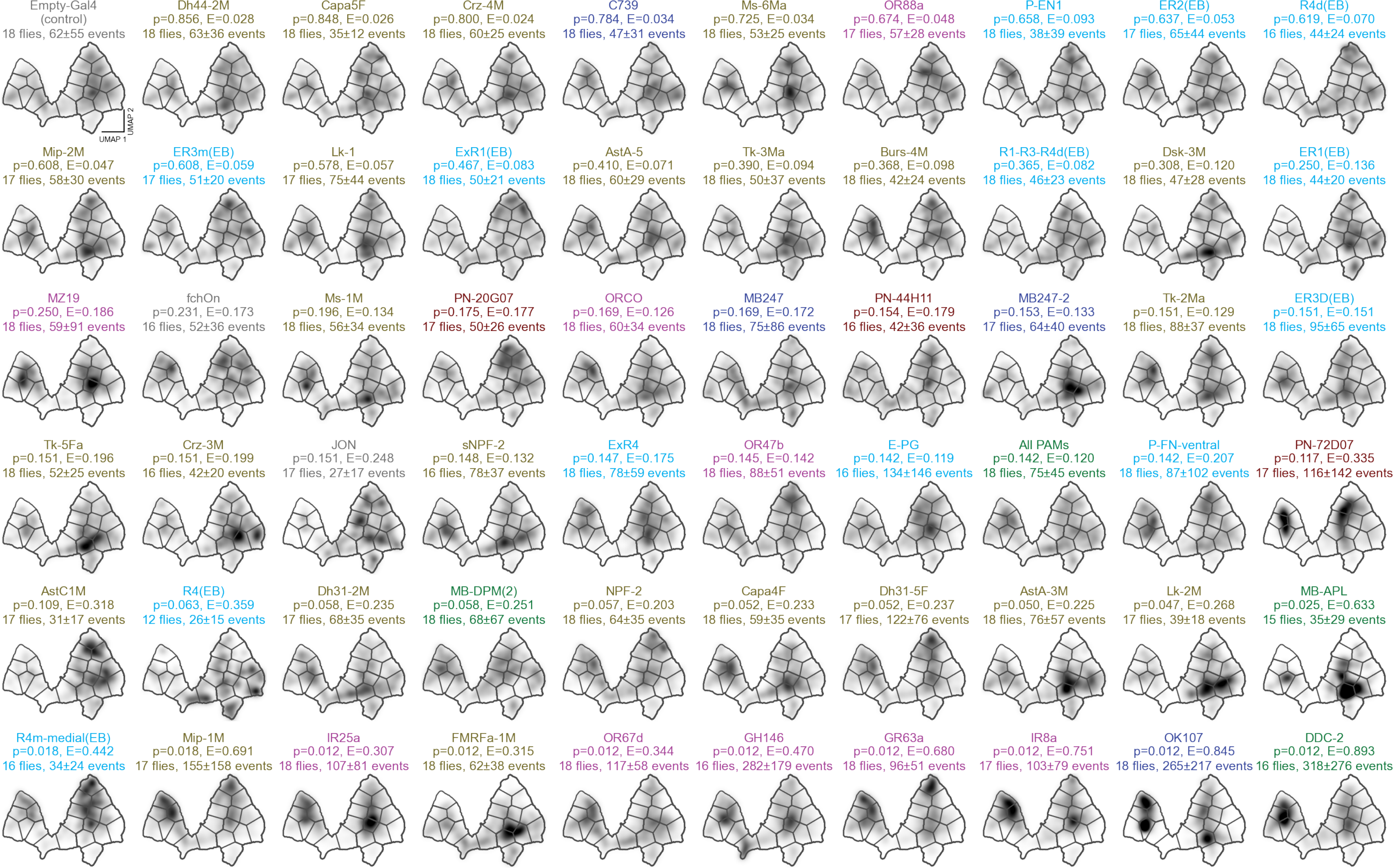

### lines_mutants.pdf

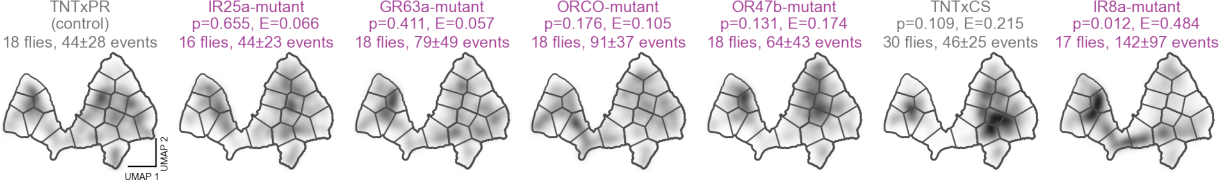

### lines_split_gal4.pdf

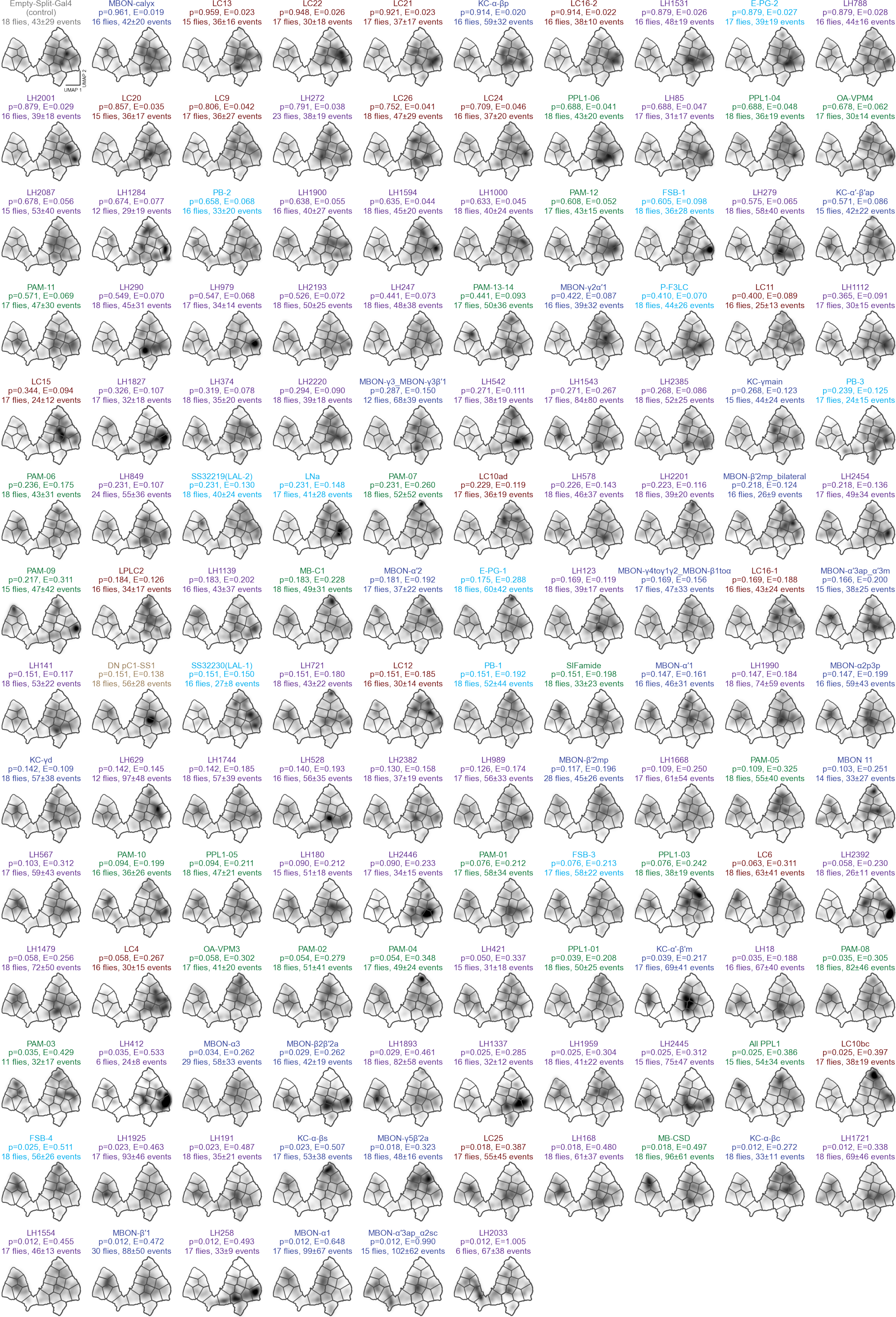

### Supplementary Information File 4

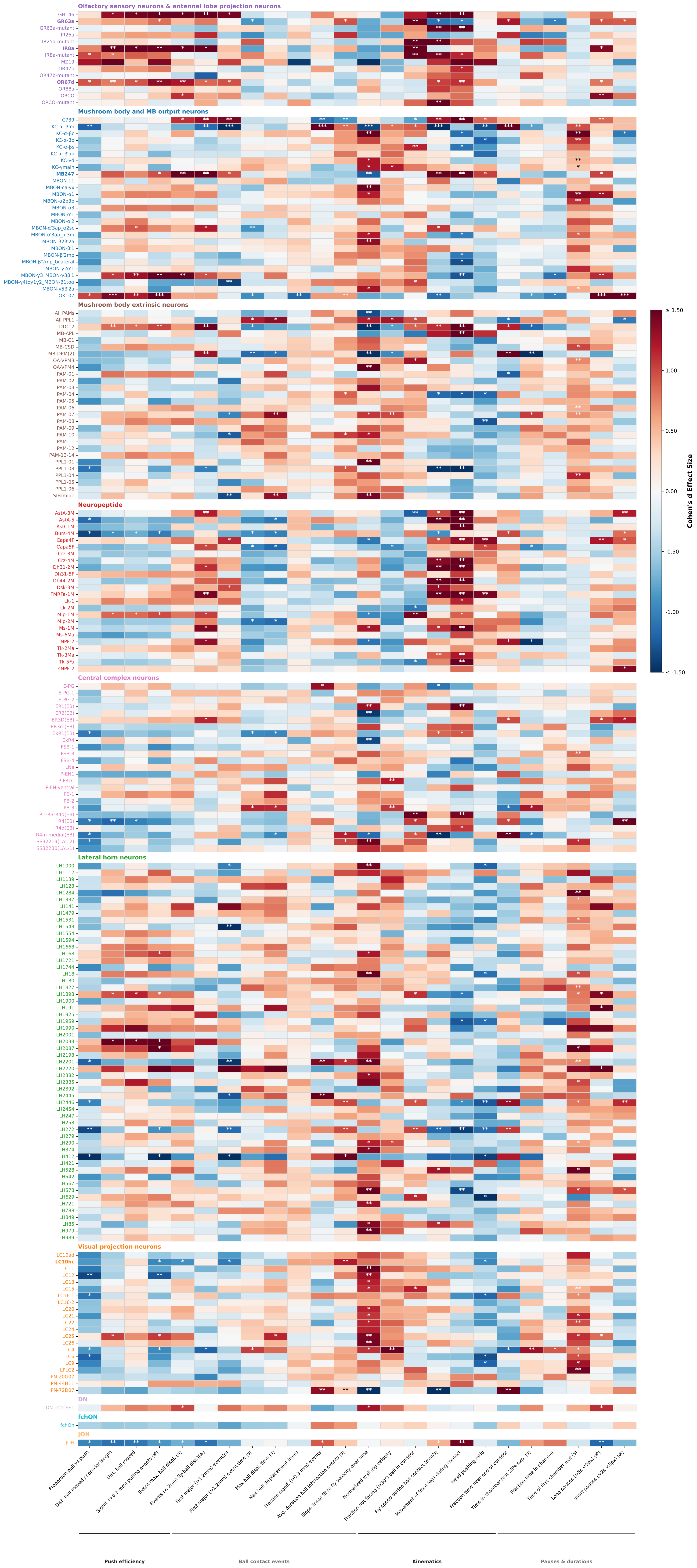

### Supplementary Information File 5

**Genotype Combined Consistency Scores**
**199 genotypes tested | 28 above 80.0% threshold**

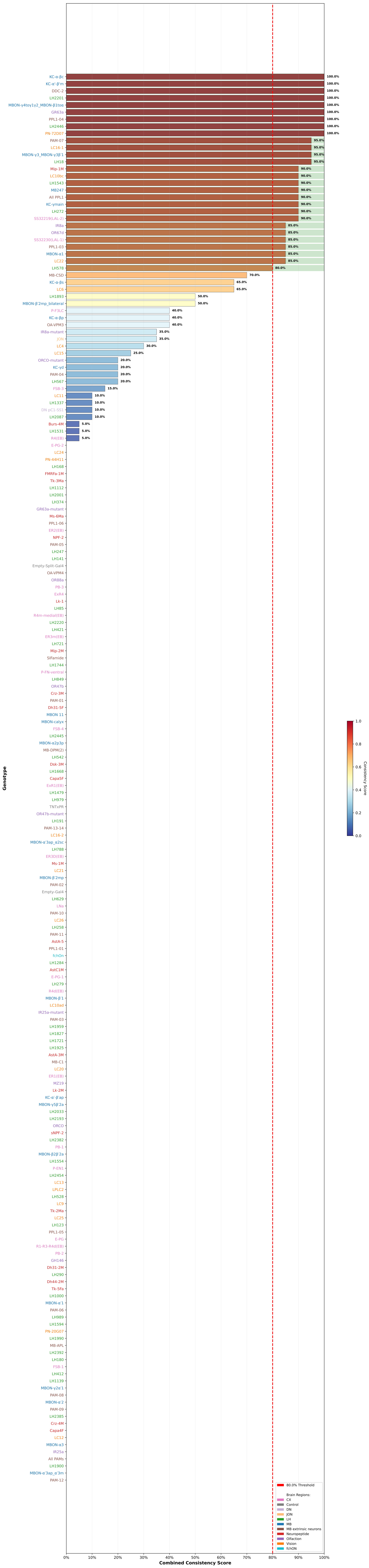
